# Actin-membrane interface stress regulates Arp2/3-branched actin density during lamellipodial protrusion

**DOI:** 10.64898/2026.03.06.710140

**Authors:** Mitchell T. Butler, Max A. Hockenberry, Harrison H. Truscott, Wesley R. Legant, James E. Bear

## Abstract

Motile cells can sense and exert forces on the extracellular environment through dynamic actin networks. Increased stress against the polymerizing barbed ends of branched actin networks has been shown to lead to an increase in the density of these networks through a force feedback mechanism, though this phenomenon has not been explored through the examination of real-time responses of endogenous branched actin in cells. Here, we utilize mouse CRISPR knock-in fibroblasts with labeled Arp2/3 complex to identify cellular and extracellular conditions that regulate branched actin density and enrichment at the leading edge of lamellipodial protrusions. A common theme shared among all branched actin density-increasing conditions is higher levels of predicted interface stress between the plasma membrane and the barbed ends of the lamellipodial actin network. Among these conditions, we find that Arp2/3 is specifically required for robust spreading and protrusion in response to increased extracellular viscosity. Interestingly, time-lapse traction force microscopy of Arp2/3-dependent viscosity responses show significantly reduced changes in strain energy applied to the substrate when compared to spreading and motility utilizing cell-matrix adhesion. In addition, we find that increased extracellular viscosity can bypass the need for extracellular matrix proteins to support lamellipodial protrusion driven by optogenetic Rac activation. Our studies provide strong support for *in vitro* models of branched actin force feedback responses and further characterize an essential role for branched actin in mediating dramatic cell shape changes in response to increased extracellular viscosity.

## INTRODUCTION

The ability of cells to sense and respond to their environment is essential for many physiological processes, such as wound healing(Shaw and Martin 2016), immune response(Friedl and Weigelin 2008, Ryan, Kim et al. 2024), neuronal pathfinding(Ayala, Shu et al. 2007), and tissue morphogenesis(Aman and Piotrowski 2010). Cells can interact with their extracellular environment through transmembrane receptors, and among these, cell adhesion receptors such as integrins connect cytoskeletal networks through cytoplasmic adaptor proteins to the extracellular matrix (ECM) to govern cell shape and behavior(Parsons, Horwitz et al. 2010, Kechagia, Ivaska et al. 2019). The linkage of ECM to the actin cytoskeleton through integrin-containing adhesions serves as a molecular clutch and allows motile cells to push and pull against the substrate, producing the traction forces that facilitate cellular translocation(Lauffenburger and Wells 2001). Importantly, cells adapt the distribution and magnitude of forces they apply on the substrate based on extracellular molecular signals and the physical properties of the environment, though there is still much to uncover regarding the hierarchy, feedback, and overlap between various mechanistic inputs.

Organization and dynamics of the actin cytoskeleton dynamics play a prominent role in the sensing and responding to the extracellular environment. Actin branches are generated through the seven-subunit Arp2/3 complex, which can bind to the side of an existing actin filaments and nucleate a new filament as a stereotyped branch with an angle of ∼70°(Mullins, Stafford et al. 1997, Svitkina and Borisy 1999). Highly dynamic branched actin networks can be found in lamellipodia, which are broad, flat cellular protrusion often oriented in the direction of cell migration. Here, the polymerization of branched actin networks against the membrane provides the force to not only push the leading cell edge forwards(Pollard and Borisy 2003), but also to cluster small groups of integrin receptors near the edge(Choi, Vicente-Manzanares et al. 2008). While most studies on the links between integrins and actin regulation have primarily focused on contractile actomyosin bundles anchored to larger and more mature integrin focal adhesions, these smaller, more dynamic nascent integrin adhesions associated with polymerizing branched actin networks also play an important role in sensing environmental cues to direct cell migration(Romero, Le Clainche et al. 2020). Dense branched actin networks have been shown to be essential for cells to sense and respond to gradients of ECM proteins(Wu, Asokan et al. 2012), and during these haptotactic responses, cell adhesion signaling promotes more persistent protrusions and migration towards higher ECM concentrations through the activity of FAK and Src family kinases(King, Asokan et al. 2016). Signaling downstream of integrin clustering upon adhesion formation leads to activation of Rac GTPase, which activates the WAVE regulatory complex (WRC) to promote the formation Arp2/3-actin branches at the leading edge of protrusions(Stradal, Rottner et al. 2004, Insall and Machesky 2009, Rotty, Wu et al. 2013, Lappalainen, Kotila et al. 2022).

In addition to molecular signaling, there is increasing evidence that physical forces can also impact Arp2/3-branched actin dynamics(Lappalainen, Kotila et al. 2022). Several *in vitro* studies have highlighted a force feedback mechanism that increases the density of branched actin networks polymerized against increased compressive forces(Bieling, Li et al. 2016, Bieling, Weichsel et al. 2022). In cells, aspiration or severing the rear of cells to manipulate membrane tension can positively or negatively influence leading edge actin density, respectively(Mueller, Szep et al. 2017), and similar observations of increased leading edge actin density have qualitatively been made following hypertonic shock(Gauthier, Fardin et al. 2011) or increased extracellular viscosity(Bera, Kiepas et al. 2022). Branched actin is also essential for cell volume control in response to osmotic stresses(Wu, Haynes et al. 2013), likely mediated through changes in actin cortex when global forces are applied to cells. However, our understanding of how forces influence branched actin has been limited by the tools available to directly observe branch formation in cells and quantitative comparisons of various mechanical inputs.

While many studies point to roles for either signaling through integrin adhesions or physical forces on Arp2/3-branched actin regulation, few studies have explored the overlap or interplay between load force and adhesion signaling and the contributions of these branched actin regulatory inputs that are shared or unique between them. We previously developed tools allowing for the optogenetic activation of endogenous Rac(Guntas, Hallett et al. 2015) and employed them in cells on a wide range of substrate plating conditions(Zimmerman, Asokan et al. 2017). From this, we found that a minimum threshold of ECM concentration must be met in order for cells to protrude in response to optogenetic Rac activation. However, we were left with several open questions, such as whether cell protrusion failed on low ECM conditions due to insufficient signaling, failed mechanical clutching of actin networks through integrins adhesions, or some combination of both. Through live imaging of the Arp2/3 complex in cells, we addressed these questions here with analyses of actin branch formation under conditions that would be predicted to increase the “load force” on barbed ends of the lamellipodial actin network pushing against the plasma membrane.

Through examination of the interplay between Arp2/3 localization, Rac signaling, extracellular conditions, and cell shape, we offer a detailed and quantitative characterization of Arp2/3-branched architecture across a range of manipulations. Cells that adopt a spread, flattened morphology upon either ECM engagement, relaxation of cortical actomyosin, or application of external physical forces all exhibit increasingly enriched branched actin at the edge of flat protrusions, and additionally, cells subjected to osmotic shock or extracellular viscosity show similar effects. Genetic loss of Arp2/3 showed that branched actin specifically is required for dramatic cell shape changes in response to increased extracellular viscosity, and in turn, increased extracellular viscosity can support both efficient cell spreading and successful protrusion following optogenetic Rac activation on low ECM conditions, shedding further light the close relationship between branched actin and mechanical forces acting on cells.

## RESULTS

### Cells with endogenously labeled Arp2/3 complex reveal striking substrate-dependent actin network architecture at the leading edge

To assess how branched actin dynamics are influenced by extracellular matrix (ECM) to control cell shape and behavior, we utilized a mouse dermal fibroblast cell line with the ARPC2 (p34) subunit of the Arp2/3 complex endogenously labeled with the mScarlet red fluorescent protein via CRISPR/Cas9 gene editing. Biallelic labeling of this subunit allows for live-cell visualization of all endogenous Arp2/3 complexes (Fig.1S1), and the genetic background of the mouse from which the parental JR20 fibroblast cell line was derived(Rotty, Wu et al. 2015) allows for conditional knockout of this same ARPC2-mScarlet tagged subunit (Table S1). To evaluate the impact of ECM engagement on Arp2/3 spatiotemporal dynamics, these fibroblasts were seeded on glass coverslips coated with either fibronectin, providing a dense ECM to engage robust integrin adhesions, or poly-L-Lysine (PLL), which promotes cell flattening and spreading through electrostatic change differences between the lysine residues and the cell surface. Although there are ECM molecules present in serum-containing media, engagement with fibronectin by cells plated on poly-L-Lysine is limited by these cells not being to produce their own fibronectin following CRISPR-induced disruption of the *Fn1* gene (Chandra, Butler et al. 2022).

Roughly one hour after plating, cells on poly-L-Lysine exhibited a broad, irregular distribution of Arp2/3, with dense accumulations surrounded by more diffuse regions along the leading edge of protrusions (Fig.1A and Movie 1). A striking difference was seen in cells plated instead on fibronectin-coated substrates, which showed a dense and more uniform enrichment of ARPC2-mScarlet at the leading edge of lamellipodia (Fig.1A and Movie 2). This ECM-dependent difference in Arp2/3 localization is generally consistent with more broad distributions of injected labeled actin and phalloidin staining shown at the edges of Swiss 3T3 fibroblasts plated on poly-L-Lysine and more narrow distributions when plated on fibronectin (Machesky and Hall 1997). Providing qualitative support for our visual impressions, line scans of ARPC2-mScarlet intensity running from protrusive cell edges inwards towards the cell body show a sharper, taller peak when cells are plated on fibronectin compared to a wider distribution with a more gradual drop-off in intensity when plated on poly-L-Lysine (Fig.1B). Measurements of the mean ARPC2-mScarlet intensity under a roughly 1 micron wide line drawn along the leading edge serves as a general measure of Arp2/3 enrichment at lamellipodial protrusions, and the values of these measurements for cells plated on fibronectin are nearly double those for cells plated on poly-L-Lysine (Fig.1C). We also compared the distributions of Arp2/3 labeling in protrusive regions by measuring the width of line scan peaks (shown in Fig.1B) at half of the maximum value, referred to as the full width half maximum (FWHM). Together, these leading edge and line scan measurements suggest that lamellipodial actin branching density is significantly higher in cells plated on fibronectin relative to being plated to poly-L-Lysine (Fig.1B-D). As would be expected with an increase in Arp2/3 density at the leading edge, the amount of free barbed ends is significantly increased in cells plated on fibronectin when compared to cells plated poly-L-Lysine (Fig.1S2A-C), confirming that the more abundant Arp2/3 complexes seen here are being incorporated into the leading edge actin network and mediating more dense actin branching.

**Figure 1.**
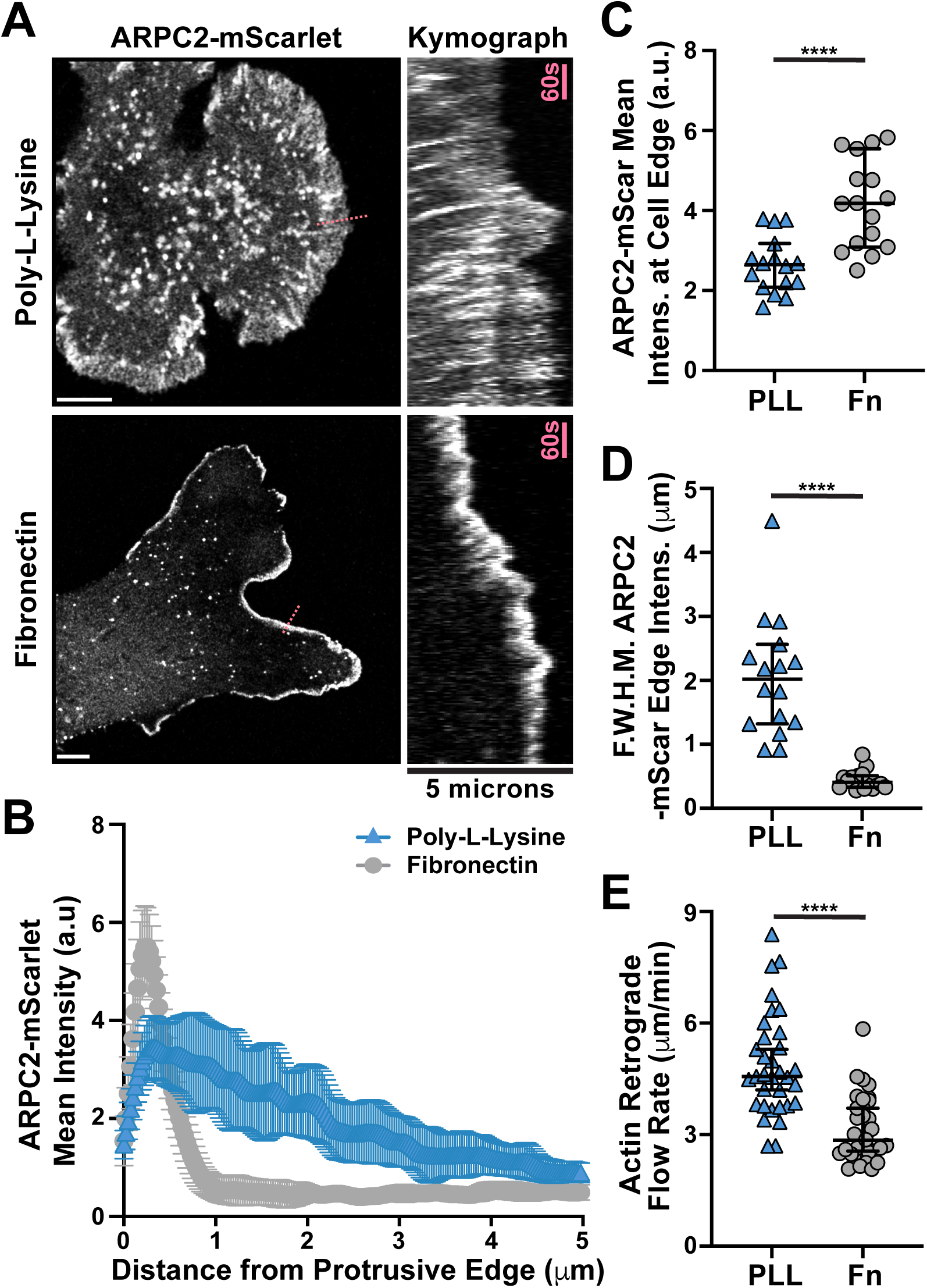
Cells with endogenously labeled Arp2/3 complex reveal striking substrate-dependent actin network architecture at the leading edge. **(A)** Frames from time lapse movies of endogenously-labeled ARPC2-mScarlet expressed by fibroblasts plated on glass coverslips coated with either fibronectin or poly-L-Lysine alongside associated kymographs of protrusive regions marked with a dashed pink line in the left panels. Scale bars span 10 microns. (B) Plot of mean ARPC2-mScarlet intensities along line scans starting near the middle of the largest cellular protrusion, aligned perpendicular to the cell edge, and directed inwards for 5 microns while avoiding regions of obvious ruffles and folds in cells plated as described in (A). n = 16 cells from 3 experiments. (C) Plot of mean ARPC2-mScarlet intensities at and within roughly 1 micron of the cell edge along the largest protrusion in fibroblasts plated as described in (A). n = 16 cells from 3 experiments. (D) Plot of width measurements in microns of ARPC2-mScarlet signal that is above 50% the maximum intensity recorded, referred to as “Full Width Half Max” and abbreviated as F.W.H.M, calculated from the line scans shown and described in (B). **(E)** Plot of retrograde flow rates visualized by bleaching GFP-β-Actin signal at the edge of protrusions in cells plated as described in (A) and measured by calculating the slope of recovering signal in kymographs generated from bleached regions. n = 33 and 32 cells for poly-L-Lysine and fibronectin conditions, respectively, from 3 experiments. Note that the data from cells on poly-L-Lysine coated glass in panels (B-E) were collected at the same time as other experimental conditions for a separate study, and values for fibronectin here have been published as the 10 μg/ml control condition previously (Chandra, Butler et al. 2022).

### Branched actin enrichment in fibroblasts plated on fibronectin depends on Integrin engagement

Early studies of GTPase signaling and cell adhesions demonstrated that Swiss 3T3 fibroblasts require extracellular matrix for the formation of integrin adhesions(Hotchin and Hall 1995), and we were able to verify that such structures are essentially absent in protrusions over poly-L-Lysine in JR20 fibroblasts through the live imaging of GFP-Paxillin (Fig.1S3A-B). Adhesion formation has been shown to coincide with a reduction in retrograde actin flow(Alexandrova, Arnold et al. 2008), and retrograde flow rates that could be considered to be rapid for fibroblasts have been observed in Xenopus XTC cells plated on poly-L-Lysine(Ryan, Petroccia et al. 2012). Thus, it seemed likely that the more diffuse and broader distribution of Arp2/3 at the leading edge of JR20 fibroblasts plated on poly-Lysine would be concomitant with an increase in retrograde flow rates over the rates seen in cells plated on fibronectin. Indeed, upon bleaching GFP-β-actin at the leading edge and monitoring the dynamic retrograde flow of subsequently polymerized F-actin, we measured significantly faster flow rates in cells plated on poly-L-Lysine relative to the dense fibronectin plating condition (Fig.1E).

To confirm the role of integrin adhesions specifically on leading edge actin organization and rule out any potential influence of alternate or secondary signaling pathways, such as an effect being mediated through Syndecans(Couchman and Woods 1999, Midwood, Valenick et al. 2004), we utilized coverslips coated with purified RGD peptides. Cells plated on glass coated with either fibronectin or RGD peptides had abundant focal adhesions and actin stress fibers, both of which were lacking in cells plated on poly-L-Lysine (Fig.1S3C). Measuring the mean ARPC2-mScarlet intensity along the very leading edge revealed that there was an increase in protrusive Arp2/3 enrichment on RGD-coated glass that was comparable to the increase in enrichment seen on fibronectin relative to poly-L-Lysine (Fig.1S3C-D). Thus, the stark differences seen in branched actin organization in cells plated on fibronectin when compared to poly-L-Lysine is likely mediated through engagement with integrin-based adhesions.

### The requirement for the Arp2/3 complex in poly-L-Lysine-based protrusions

The observation of leading-edge Arp2/3 in protrusions over surfaces coated with poly-L-Lysine suggests the presence of nucleation promoting factors (NPFs) to initiate branches(Rotty, Wu et al. 2013) and branch-promoting factors, such as Cortactin(Weed, Karginov et al. 2000, Weaver, Karginov et al. 2001), to stabilize them. These factors and their roles in branched actin organization have been well characterized(Gautreau, Fregoso et al. 2022), but the influence of ECM on their localization and behavior, including direct comparison between different substrates, have not be studied in great detail. Although we would not predict branch-promoting factors being absent here, there is some evidence that might suggest that the lack of dense Arp2/3-branched actin observed on poly-L-Lysine could be due to reduced Arp2/3-branch generation or insufficient branch stabilization. For example, integrin adhesion signaling mediated through focal adhesion kinase (FAK or Ptk2) and Src of the Src family kinases has been shown to promote WAVE NPF activity(Romero, Le Clainche et al. 2020). In addition, Cortactin has been shown to be a target for FAK(Tomar, Lawson et al. 2012) and phosphorylated by Src(Wu and Parsons 1993), though it is less well understood how these kinases influence Cortactin localization and activity.

To examine the role of actin branch regulators in the ECM-dependent phenotypes we observed, we next assessed the localization of WAVE1 (NPF) and Cortactin when cells were plated in the absence of dense ECM. Regardless of whether cells were plated on poly-L-Lysine or fibronectin, there was abundant WAVE1 at the cell periphery (Fig.2A-B and Movies 3-4), and Cortactin localized throughout the branched actin network during cell protrusion, exhibiting similar localization and dynamics to labeled Arp2/3 on both substrates (Fig.2C-D). We also tested the functional requirement for Arp2/3 in producing the broad lamellipodia-like protrusions observed on poly-L-Lysine. To accomplish this, we induced Cre-mediated recombination at the conditional *Arpc2* locus and plated cells on poly-L-Lysine. When plated on poly-L-Lysine, control cells with Arp2/3 intact displayed the expected wide, cohesive protrusions with sparse Arp2/3 branching, but Arp2/3 null cells displayed only narrow bundled actin protrusions, which appear as separate structures lacking the more diffusively labeled meshwork between the bundled actin structures (Fig.2E). We conclude that differences seen in branched actin density between poly-L-Lysine and fibronectin are not due to an inability for branches to be nucleated or stabilized on poly-L-Lysine, and instead, could be due to differences in levels of activity for NPFs or Rac GTPase signaling and/or mechanical forces influencing actin network organization.

**Figure 2.**
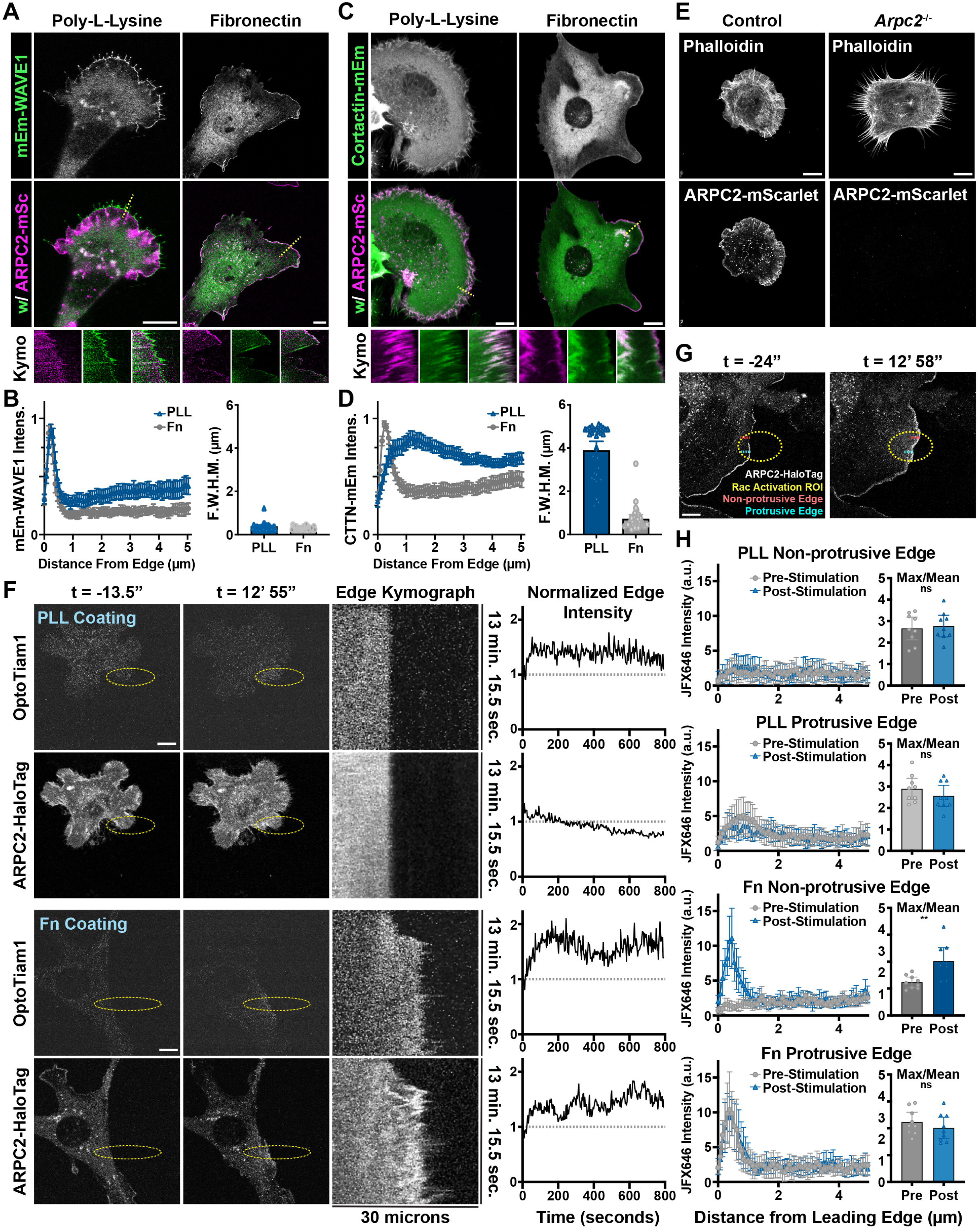
Increased Rac activation is not sufficient to induce dense lamellipodial branched actin organization in cells plated on poly-L-Lysine. **(A)** Frames from time lapse movies of endogenous ARPC2-mScarlet and exogenous mEmerald-WAVE1 stably expressed by fibroblasts plated on glass coverslips coated with either fibronectin or poly-L-Lysine and associated kymographs of protrusive regions marked with a dashed red line. Scale bars span 10 microns. **(B)** Line scans of mEmerald-WAVE1 intensity running from the leading edge of protrusions to 5 microns inward towards the cell body and Full Width Half Max measurements of the width, in microns, of the line scan peaks at half their maximum value. n = 30 cells for each plating condition from two separate experiments. **(C**) Frames from time lapse movies of endogenously ARPC2-mScarlet and exogenous Cortactin-mEmerald stably expressed by fibroblasts plated on glass coverslips coated with either fibronectin or poly-L-Lysine and associated kymographs of protrusive regions marked with a dashed red line. Scale bars span 10 microns. **(D)** Line scans of Cortactin-mEmerald intensity running from the leading edge of protrusions to 5 microns inward towards the cell body and Full Width Half Max measurements of the width, in microns, of the line scan peaks at half their maximum value. n = 32 cells for each plating condition from two separate experiments. **(E)** Images of control JR20 fibroblasts with Arp2/3 intact and 4HT-treated *Arpc2* knockout fibroblasts plated on poly-L-Lysine coated surfaces and visualized using fluorescently labelled phalloidin. Scale bars span 10 microns. **(F)** Images from timelapse movies of Tiam1-DH/PH-TagRFPt-SspBmicro and ARPC2-HaloTag (labeled with JF646 Halo Ligand) stably expressed by 4HT-treated *Arpc2* knockout fibroblasts that have been plated on glass coverslips coated with poly-L-Lysine (top panels) or fibronectin (bottom panels) and regularly stimulated with 405 nm light inside the region of interest (ROI) labeled with a yellow dashed oval starting at t = 0. These cells additionally express Venus-iLid-CAAX (not shown). The associated kymograph was generated from a line (not shown) bisecting the yellow dashed oval ROI along the long axis, and the associated plot is a measure of the intensity of the associated marker within ∼1 micron of the cell edge down the length of the accompanying kymograph. Scale bars span 10 microns. **(G)** Images from a timelapse movies taken before and after optogenetic stimulation of endogenous Rac showing the distribution of ARPC2-HaloTag labeled with JF646 Halo ligand. The ROI used for optogenetic stimulation is shown as a yellow dashed oval and overlapped regions of the cell that were protrusive (Halo signal enriched) and non-protrusive (strong Halo signal absent) pre-stimulation. Red and cyan dashed lines mark representative regions of the cell from which line scan intensities were collected, shown in (H). **(H)** Plots of values from line scan intensity measurements running from the edge of the cell to 5 microns inwards toward the cell body taken prior to and following optogenetic stimulation of endogenous Rac as well as plots of the maximum/mean values of individual line scans. Fibroblasts were plated on glass coverslips coated with either poly-L-Lysine or fibronectin, and the subregions of the cell periphery measured were within the stimulation ROI and either labeled by ARPC2-HaloTag and thus protrusive or not labeled and considered to be non-protrusive prior to optogenetic stimulation. Measurements were also taken in equivalent region following roughly 13 minutes of stimulation as shown in (G) for comparison. n = 9 cells for each plating condition from two separate experiments.

### Increased Rac activation is not sufficient to induce dense branched actin organization on poly-L-Lysine

Although cells plated on poly-L-Lysine exhibit Arp2/3-activating NPFs (such as WAVE) at the leading edge of protrusions, one possible explanation for the sparse density of branched actin observed under these conditions is insufficient levels of integrin-mediated signaling that activate the NPFs localized here. Signaling from nascent integrin adhesions at the leading edge activates kinases such as FAK and Src, leading to Rac activation and cell protrusion(Tilghman, Slack-Davis et al. 2005, Owen, Pixley et al. 2007, Choi, Zareno et al. 2011, King, Asokan et al. 2016). Similarly to previous reports(Hotchin and Hall 1995), nascent adhesions are markedly reduced or absent in JR20 protrusions over poly-L-Lysine (Fig.1S3), and we would expect that levels of any protrusion-promoting signaling would be low under these conditions. We previously employed an iLid-based optogenetic system to concentrate the DHPH domain of the RacGEF Tiam1 at the cell membrane and activate endogenous Rac, and doing so is sufficient to drive cell protrusion in fibroblasts plated on fibronectin, but not on poly-L-Lysine (Zimmerman, Asokan et al. 2017). Despite Rac activation in cells plated on poly-L-Lysine not being sufficient to drive cell protrusion, we did not previously examine whether it could drive branched actin enrichment at the leading edge, independent of protrusive behavior.

To test whether Rac activation alone was sufficient to induce the formation of dense leading-edge Arp2/3-branched actin in cells plated on poly-L-Lysine, we generated a *Arpc2* knockout, ARPC2-HaloTag rescue cell line to facilitate visualization of Arp2/3 complexes simultaneously with an iLid-based optogenetic system for activation of endogenous Rac we utilized previously (Zimmerman, Asokan et al. 2017). Consistent with our previous findings, we found that while optogenetic Rac activation increases lamellipodial ruffling, it does not drive robust cell protrusion on poly-L-Lysine, and additionally, we observed negligible changes in the distribution of labeled Arp2/3 complexes in stimulated regions (Fig.2F and Movie 5). In contrast, control cells extended robust cell protrusions over fibronectin driven by optogenetic Rac activation and exhibited dense branched actin within such protrusions (Fig.2F and Movie 6). We repeated similar Rac activations in cells plated on poly-L-Lysine and fibronectin using optogenetic stimulation ROIs that overlapped both protrusive regions that had Arp2/3 strongly localized there and non-protrusive regions that showed no obvious Arp2/3 localization. Line scan intensity measurements of labeled Arp2/3 at protrusive and non-protrusive cell edges prior to and following Rac activation (Fig.2G) revealed changes in branched actin distribution only when cells are plated on fibronectin and only in regions that are non-protrusive prior to activation (Fig.2H). These results indicate that increased Rac activity is insufficient for dense actin branching in cell protrusions, and since activation of Rac has been identified as one of the key signaling events downstream of integrin signaling to support cell protrusion, this result also suggests that differences in integrin-mediated signaling alone are unlikely to account for the differences in Arp2/3 distribution in cells plated on the different substrates.

### Enrichment of Arp2/3-branched actin is linked to cell spreading

To begin to assess how cell shape and mechanics might influence the branched actin differences seen between fibronectin and poly-L-Lysine plating conditions, we imaged JR20 fibroblasts using scanning electron microscopy (SEM). Cells plated for roughly forty-five minutes on fibronectin showed completely flat, smooth protrusions, but cells plated on poly-L-Lysine showed abundant ruffles, filopodia, and folds in the plasma membrane (Fig.3S1A-B). Such structures seen in cells plated on poly-L-Lysine can act as “reservoirs” for excess plasma membrane by having a high cell membrane surface area to cell volume ratio and may indicate lower plasma membrane tension in cells plated on poly-L-Lysine. Furthermore, it also suggests that cells may be spreading at very different rates under these conditions, and measuring spread area over time confirmed this notion (Fig.3S1C). In extended time-lapse movies of cells plated on poly-L-Lysine, we observed enrichment of the Arp2/3-branched actin when the cells reached what appeared to be their maximum spread area (Fig.3S1D). Interestingly, some cells plated on poly-L-Lysine would spread out transiently and then shrink back to a smaller footprint, and the Arp2/3-branched actin density would appear maximally enriched during the period of maximum spreading and appear less enriched afterwards when spread area was subsequently reduced (Fig.3S1D and Movie 7).

To quantitatively test the idea that Arp2/3-branched actin density was linked to spread area, we examined branched actin content relative to cell spread area over time in cells plated on either poly-L-Lysine (Movie 7) or fibronectin (Movie 8). For this characterization, we used live-cell imaging to record the entire spreading process and generated ARPC2-mScarlet intensity profiles around the cell periphery at each time point. We related these values of cell spread area to the maximum area observed (Fig.3A & Fig.3S2). From these profiles, we calculated FWHM values that show cells plated on fibronectin tend to have more enriched branched actin throughout the spreading process relative to poly-L-Lysine plating, but cells plated on either substrate have branched actin that is increasingly confined to the outermost edge of cell protrusions as cells continue to spread (Fig.3B). The ratio of the maximum intensity over the mean intensity throughout the cell periphery at each measured time point is also a useful metric for Arp2/3 density at the edge of cells (Fig.3S2), and these values similarly show that branched actin becomes increasingly enriched in protrusions throughout the spreading process and that cells plated on fibronectin produce denser branched actin networks sooner, possibility due to more robust clutching of retrograde actin network flow (Fig.3C-D). Importantly, we recorded very similar minimum and maximum projected cell spread areas for both substrates (Fig.3E), indicating the calculated Arp2/3 distributions were for cells adopting similar, nearly fully spread states under both conditions.

**Figure 3.**
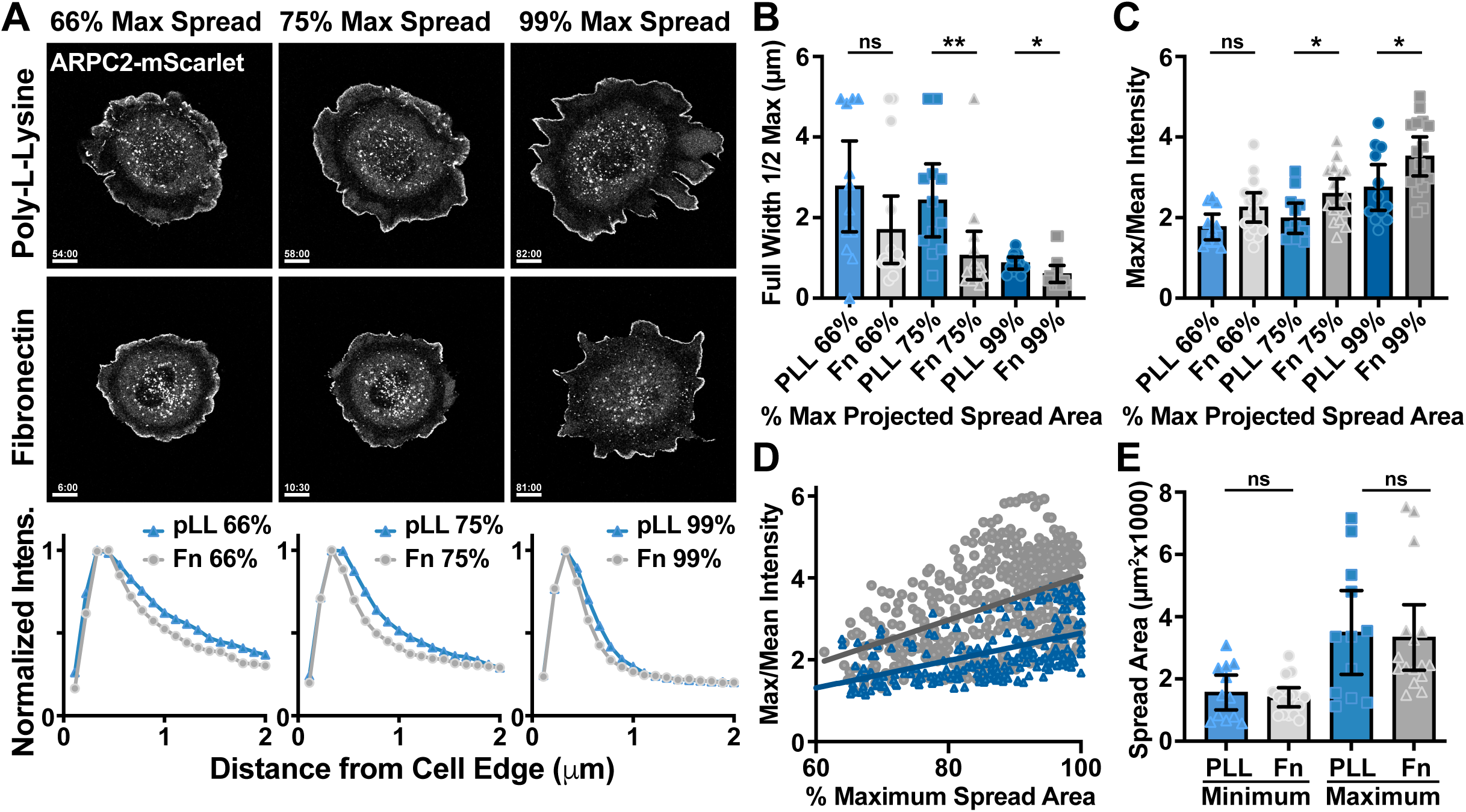
Enrichment of Arp2/3-branched actin is linked to cell spreading. **(A)** Representative images of ARPC2-mScarlet endogenously expressed by fibroblasts plated on glass coverslips coated with either poly-L-Lysine or fibronectin as they reached a projected cell spread area of 66%, 75%, and 99% of the maximum spread area recorded. Graphs below the image panels show the distribution of intensity around the cell periphery at a given distance from the cell edge for each of the two plating conditions shown above. Scale bar spans 10 microns. **(B)** Plot of Full Width Half Max ARPC2-mScarlet mean intensity around the cell periphery at various stages of spreading relative to the maximum recorded area when plated on surfaces coated with poly-L-Lysine or fibronectin. **(C)** Plot of the maximum value from a range of mean ARPC2-mScarlet intensities measured in 1-pixel step intervals around the cell spanning from the cell edge to 5 microns inward, which is divided by the mean value of all intensities throughout this same range and shown as an alternative measure for the values shown in (B). Quantification of these values is graphically detailed in Figure 3 Supplement 1 for clarity. **(D)** Plot of the maximum/mean ARPC2-mScarlet intensity within 5 microns of the cell edge around the entirety of the cell periphery against cell spread area as a percentage of the maximum recorded, showing a significantly steeper slope for these values when measured in groups of cells plated on fibronectin compared to those plated on poly-L-Lysine. **(E)** Minimum and maximum projected spread areas of the cells measured. n = 12 for poly-L-Lysine and 16 for fibronectin from 3 experiments for (B-E).

### Inducing spreading through Myosin inhibition promotes enrichment of branched actin in cell protrusions

Slow spreading of cells plated on poly-L-Lysine leads to a gradual enrichment of branched actin at the cell periphery (Fig.3), and we next wanted to assess the effects of inducing rapid cell spreading on poly-L-Lysine with a similar timescale as adhesion-based spreading seen on surfaces coated with dense fibronectin ECM. Addition of para-aminoblebbistatin inhibits non-muscle myosin II and would be predicted promote cells to more readily maximize their surface contact area with the electrostatically attractive poly-L-Lysine surface coating when flattening is no longer resisted by the cortical actomyosin contractility(Cai, Biais et al. 2006). With some similarities to previous reports that examined the effect of myosin activity levels on cell spreading on surfaces coated with fibronectin or poly-L-Lysine(Flinn and Ridley 1996, Cai, Rossier et al. 2010, Gauthier, Fardin et al. 2011), treatment of cells plated on poly-L-Lysine with 50 μM para-aminoblebbistatin led to a rapid increase in spread cell area, with around a 2-fold increase on average relative to DMSO-treated controls (Fig.4A-B and Movies 9-10).

**Figure 4.**
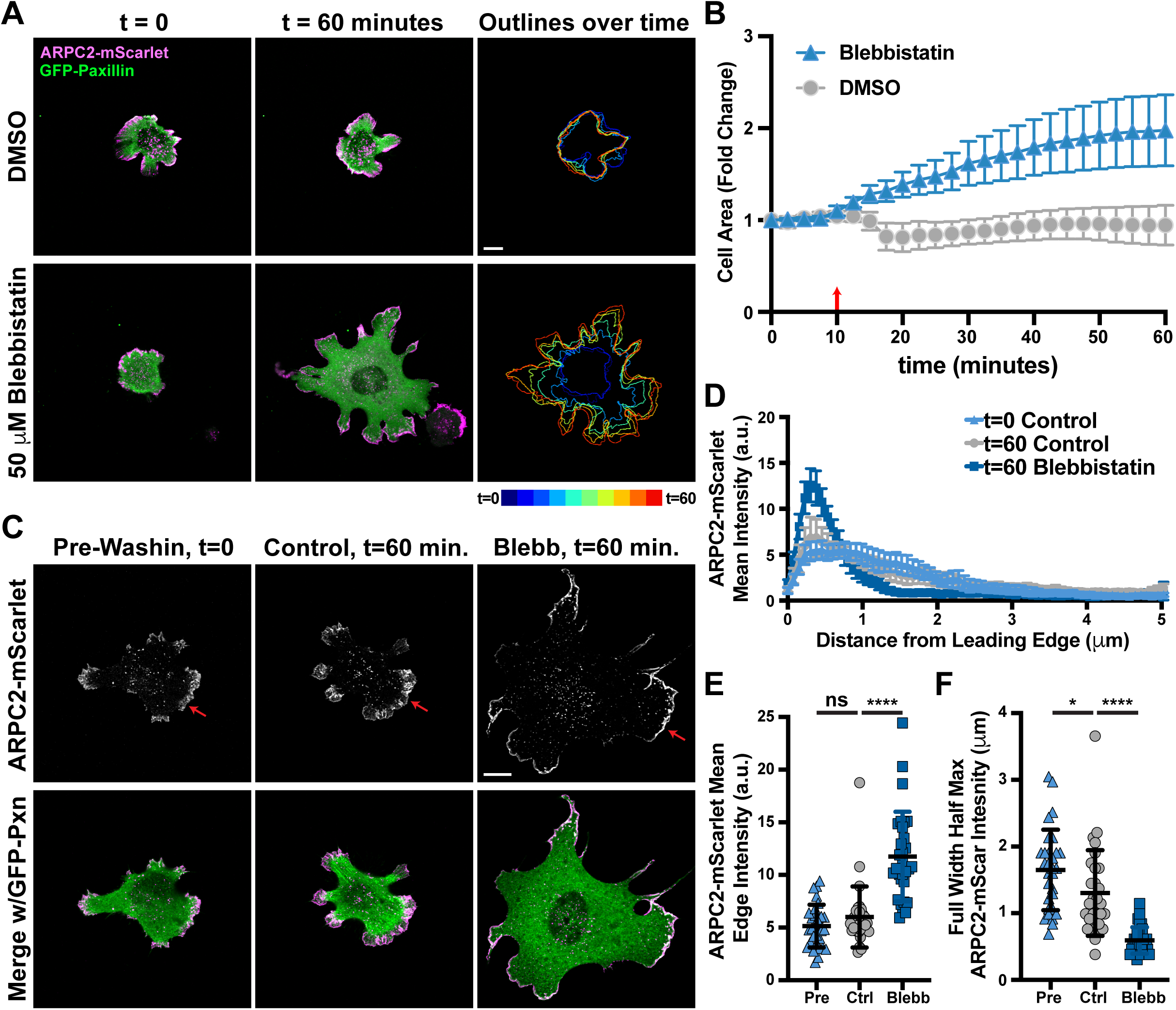
Inducing cell spreading on poly-L-Lysine with myosin inhibition results in enrichment of branched actin in cell protrusions. **(A)** Frames from representative time lapse movies of endogenous ARPC2-mScarlet and exogenous GFP-Paxillin stably expressed by fibroblasts plated on glass coverslips coated with poly-L-Lysine and treated with either 50 μM para-aminoblebbistatin (Blebbistatin or Blebb) or DMSO control at t = 10 minutes showing the change in projected cell spread area over time. Scale bar spans 10 microns. **(B)** Plot of projected cell spread area over time measured among cells shown and described in (A), with the red arrow at t = 10 minutes marking when blebbistatin was added to the media. n = 25 from 2 experiments for 50 μM para-aminoblebbistatin treatment and 14 for DMSO-treated cells. **(C)** Representative images of cells similarly treated as described in (A) and fixed either 10 minutes prior to blebbistatin addition or 50 minutes after addition for accurate measurements of ARPC2-mScarlet distribution. Scale bar spans 10 microns. **(D)** Line scans of ARPC2-mScarlet intensity along lines 5 microns long and running inward from the cell edge at the largest protrusion under the indicated conditions. n = 30 cells from 2 experiments for all three groups. **(E)** Plot of mean ARPC2-mScarlet intensities at and within roughly 1 micron of the cell edge along the largest protrusion from the same cells measured for (D). **(F)** Plot of Full Width Half Max values calculated from the line scans shown in (D).

To measure the effects of more rapid cell spreading on poly-L-Lysine on branched actin organization, para-aminoblebbistatin treatments of cells plated on poly-L-Lysine were repeated while only capturing images at the first and final time points to mitigate the effects of photobleaching (Fig.4C). The distribution of Arp2/3 in protrusions at initial timepoints one hour post-plating appeared very similar to as what was seen in cells plated on poly-L-Lysine for one hour previously (Fig.1B), with shorter, shallow peaks in lines scans (Fig.4D), relatively lower intensities along the leading edge (Fig.4E), and broader distribution of signal back from the leading edge (Fig.3F). When cells were treated with DMSO at the one-hour timepoint and examined one hour later, they showed little change in branched actin distribution, but in contrast, cells plated on poly-L-Lysine and treated with para-aminoblebbistatin showed branched actin distributions very similar to cells plated on fibronectin (Fig.1B), with tall, narrow peaks in line scans (Fig.4D), higher mean intensities along the edge of protrusions (Fig.4E), and much more narrow distribution of ARPC2-mScarlet signal confined very close to the leading edge (Fig.4F). These results suggest that forcing cells to adopt a fully spread and flattened shape is sufficient to promote dense Arp2/3-branched actin at the leading edge of protrusions. However, we could not rule out other potential effects of myosin inhibition other than promoting cell spreading, so we sought to examine how multiple other treatments to manipulate cell shape affect Arp2/3 distribution.

### Physically flattening cells or manipulating membrane tension through osmotic pressure leads to enriched protrusive branched actin

Because relieving contractility with para-aminoblebbistatin treatment can have secondary effects, such as potentially influencing F-actin turnover(Medeiros, Burnette et al. 2006, Yamashiro, Tanaka et al. 2018), we next attempted to force cells to adopt a similar spread, flattened shape while leaving non-muscle myosin II activity intact. Physically flattening cells with weighted agarose pucks on poly-L-Lysine forced cells to spread in a rapid and robust manner (Fig.4S1A-B and Movie 11). Upon physical compression, line scans demonstrate an obvious shift in the distribution of Arp2/3 to be more enriched towards the leading edge (Fig.4S1C). These finding provide additional support that cells adopting a spread, flattened shape can strongly influence branched actin density in cell protrusions.

As both para-aminoblebbistatin treatment and physical compression result in not only increased plasma membrane tension as cells spread out, but also flatter, more spatially confined cellular protrusions, we next wanted to determine whether we could manipulate these variables independently. We predicted that manipulation of cell volume through osmotic pressure could potentially be used to elicit flattening of protrusions without dramatically affecting cell spread area. When cells are plated on poly-L-Lysine, addition of 0.25M sorbitol to the medium introduces hyperosmotic conditions and initially caused a reduction in cell volume and spread cell area, but cells were then able to adapt and return to their original spread area roughly 30 min later and continue to spread slightly beyond their original projected area (Fig.4S1D-E and Movie 12). Despite the reduction in rate and magnitude of spreading seen, changes in the distribution of branched actin at the leading edge was very similar to both para-aminoblebbistatin and compressive treatments, with line scans and measurements of Arp2/3 density and enrichment closely resembling those seen when plating cells on fibronectin-coated substrates (Fig.4S1F-H). Interestingly, while SEM imaging reveals smooth, flattened cellular surfaces following one hour of para-aminoblebbistatin treatment, cells treated with 0.25M sorbitol typically retained abundant filopodial as well as membrane ruffles and folds seen in cells plated on poly-L-Lysine without additional treatments (Fig.4S1I-J). Although osmotic shock can induce a variety of cell signaling and behavior responses(Hoffmann, Lambert et al. 2009, Guo, Pegoraro et al. 2017), this serves as an additional example of experimental manipulations that we predict would increase resistance to cell protrusion and that we show leads to actin networks with more dense Arp2/3 branching.

### Increased extracellular viscosity can induce cell spreading in the absence of dense ECM and promotes enriched actin network branching in lamellipodial protrusions

Relieving cortical contractility, physically flattening cells, and manipulations of volume all similarly lead to dramatic increases in branched actin density in cell protrusions, but they all potentially impact the cell in ways that could have secondary effects on the actin cytoskeleton independent of increasing actin membrane interface stress. To supplement these finding with an additional approach, we chose to increase the viscosity of the media, which has been shown to promote increased cell motility (Bera, Kiepas et al. 2022) and cell spreading on fibronectin through a mechanism that seemingly involves suppression of lamellipodial ruffling(Pittman, Iu et al. 2022). Adding methylcellulose of the 1500 cP grade to the media at a final concentration of 0.6% increases the extracellular viscosity nearly 8-fold, from roughly 1 cP to 8 cP, and similar increases have been shown to not have a significant effect on the osmolarity of the media(Bera, Kiepas et al. 2022). This addition of 0.6% methylcellulose elicited a very dramatic spreading response in cells plated on poly-L-Lysine, resulting in a nearly 2-fold increase in projected cell spread area while untreated controls showed little changes (Fig.5A-B and Movies 13-14). Interestingly, the cell spreading response upon increase in extracellular viscosity was similar in both magnitude and rate when compared to para-aminoblebbistatin treatment despite both inducing spreading responses through what are likely very different mechanisms.

**Figure 5.**
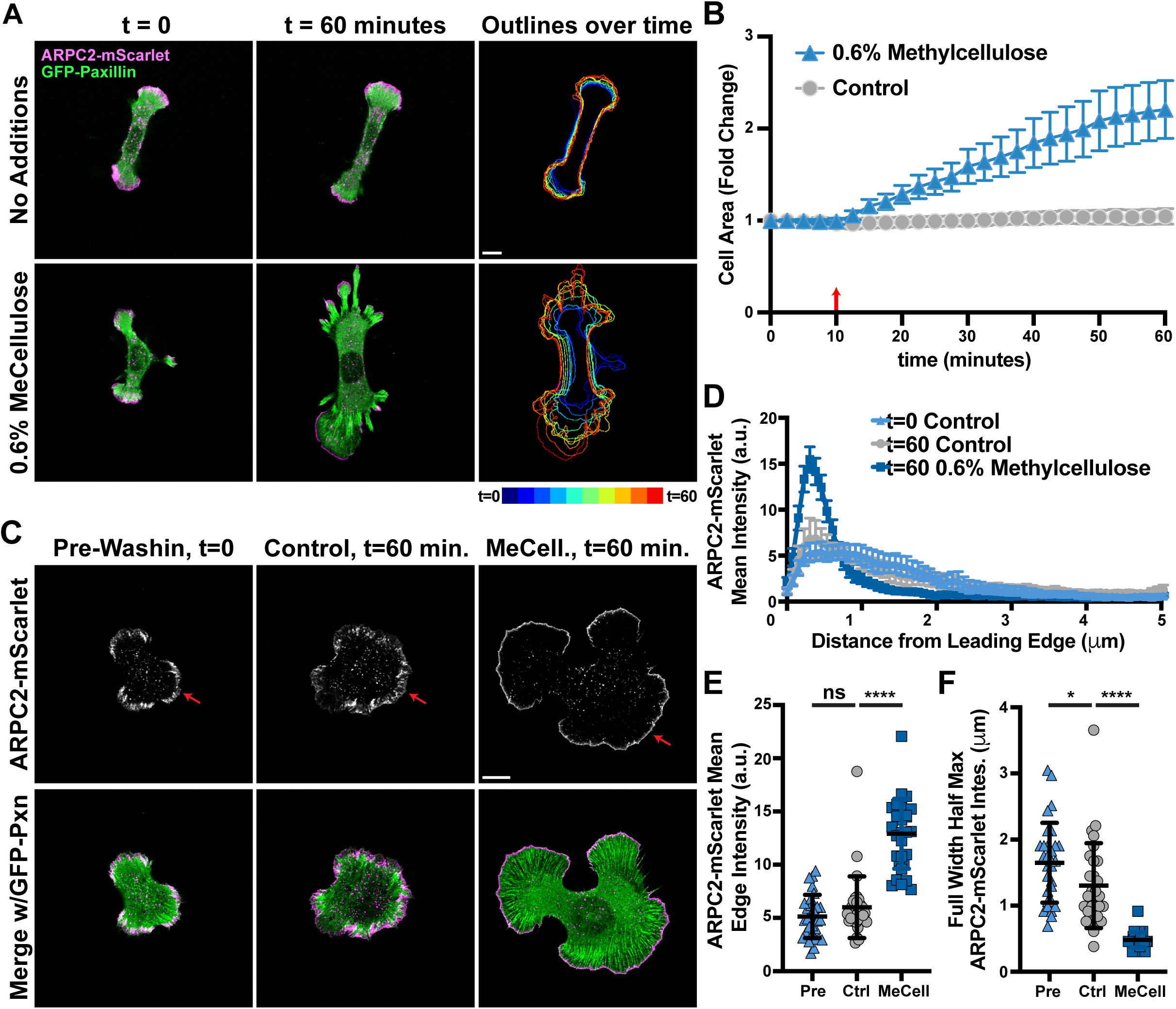
Increased extracellular viscosity can induce cell spreading in the absence of dense ECM and promotes enriched actin network branching in lamellipodial protrusions. **(A)** Frames from representative time lapse movies of endogenous ARPC2-mScarlet and exogenous GFP-Paxillin stably expressed by fibroblasts plated on glass coverslips coated with poly-L-Lysine and either treated with 0.6% methylcellulose at t = 10 minutes or left untreated, showing the change in projected cell spread area over time. Scale bar spans 10 microns **(B)** Plot of projected cell spread area over time measured among cells shown and described in (A), with the red arrow at t = 10 minutes marking when 0.6% methylcellulose was added to the media. n = 44 from 2 experiments for each condition. **(C)** Images of cells similarly treated as described in (A) and fixed either 10 minutes prior to methylcellulose addition or 50 minutes after addition for accurate measurements of ARPC2-mScarlet distribution. Scale bar spans 10 microns. **(D)** Line scans of ARPC2-mScarlet intensity along lines 5 microns long and running inward from the cell edge at the largest protrusion under the indicated conditions. n = 30 cells from 2 experiments for all three groups. **(E)** Plot of mean ARPC2-mScarlet intensities at and within roughly 1 micron of the cell edge along the largest protrusion from the same cells measured for (D). **(F)** Plot of Full Width Half Max values calculated from the line scans shown in (D).

Similar to the cell spreading events we previously examined (Figs.2-4), increasing extracellular viscosity appeared to result in an enrichment of branched actin at leading edges during time lapse imaging (Fig.5A). Images captured both ten minutes before and fifty minutes after methylcellulose allowed for accurate measurement of the effects of extracellular viscosity on the organization of the branched actin network (Fig.5C). When cells are plated on poly-L-Lysine and induced to spread with methylcellulose addition, the distribution of endogenously labeled Arp2/3 shows taller, narrower peaks in line scans (Fig.5D), higher mean intensities along the edge of protrusions (Fig.4E), and more confined distribution of leading-edge ARPC2-mScarlet signal (Fig.5F) in comparison to untreated controls. Together, these data show that increased extracellular viscosity can induce cell spreading on surfaces coated with poly-L-Lysine and consequently increase the density of the Arp2/3-branched actin network at the leading edge.

### Robust spreading in response to increased extracellular viscosity requires Arp2/3-branched actin

Some cellular responses to extracellular viscosity have been shown to involve actin branching through the use of Arp2/3 pharmacological inhibitors. As we found a dramatic increase in endogenous branched actin density in cells spreading on poly-L-Lysine upon increasing extracellular viscosity, we set out to assess the role of branched actin in this spreading response using our conditional *Arpc2* genetic knockout. Upon loss of Arp2/3 through Cre-mediated recombination at the *Arpc2* locus, we did not observe any spreading of the cells in response to 0.6% methylcellulose treatment (Fig.6A-B). In addition, neither wild type nor Arp2/3 null cells showed any change in cell area upon treatment with methylcellulose when already spread out on fibronectin (Fig.6S1). However, loss of Arp2/3 did not prevent cell spreading on poly-L-Lysine in response to para-aminoblebbistatin treatment (Fig.6C-D), likely due to differences in spreading mechanisms between methylcellulose and blebbistatin treatments. These observations confirm that branched actin is required for robust cell spreading in response to increased extracellular viscosity conditions.

**Figure 6.**
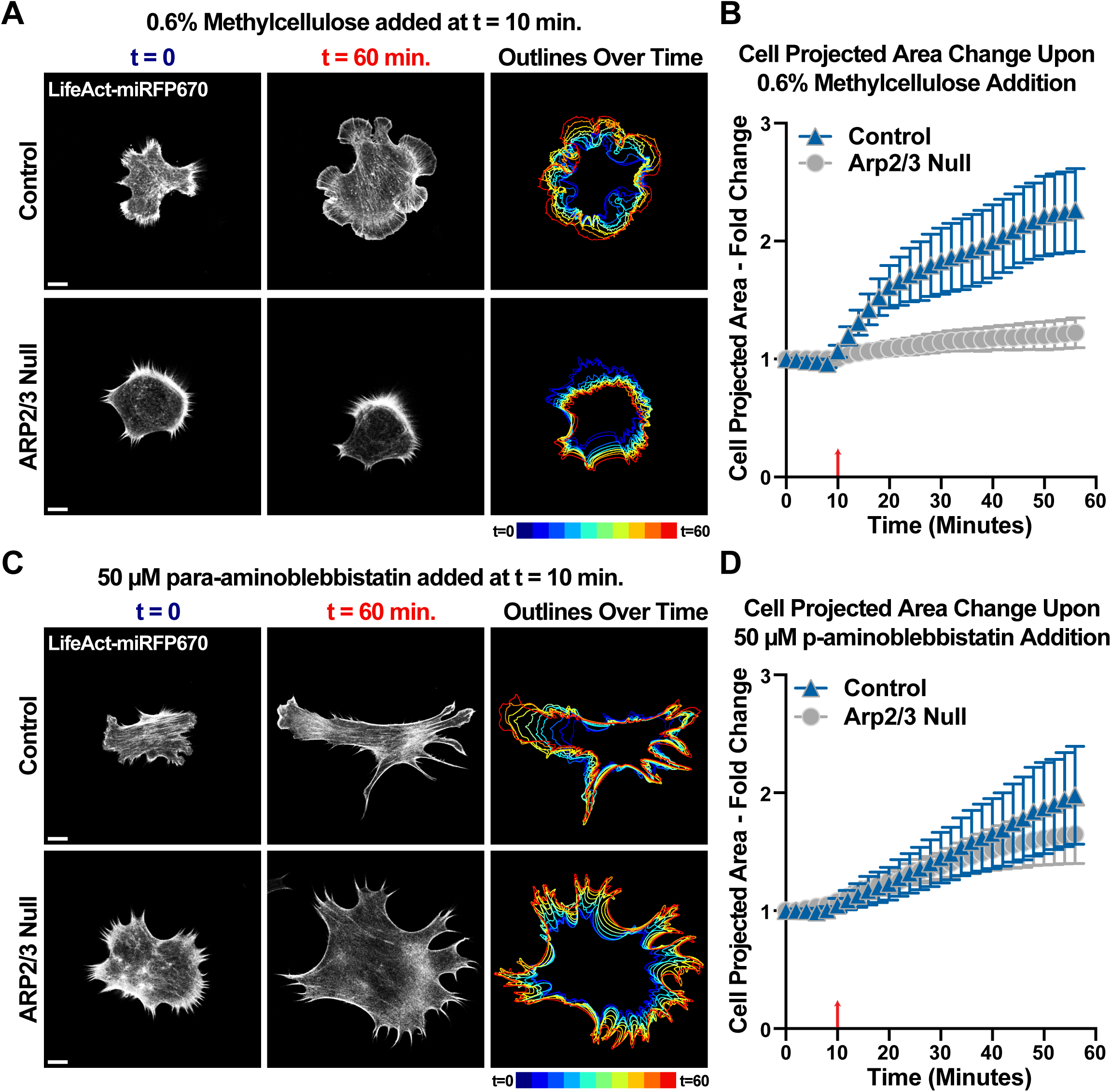
Robust spreading in response to increased extracellular viscosity requires Arp2/3-branched actin. **(A)** Frames from representative time lapse movies LifeAct-miRFP670 stably expressed by 4-HT-treated *Arpc2* knockout fibroblasts and control cells with Arp2/3 intact plated on glass coverslips coated with poly-L-Lysine and treated with 50 μM para-aminoblebbistatin at t = 10 minutes. Scale bars span 10 microns. **(B)** Plot of projected cell spread area over time measured among cells shown and described in (A), with the red arrow at t = 10 minutes marking when 0.6% methylcellulose was added to the media. n = 27 for control and 28 for Arp2/3 null cells from 2 experiments. **(C)** Frames from representative time lapse movies of LifeAct-miRFP670 stably expressed by 4-HT-treated *Arpc2* knockout fibroblasts and control cells with Arp2/3 intact plated on glass coverslips coated with poly-L-Lysine and treated with 0.6% methylcellulose at t = 10 minutes. Scale bars span 10 microns. **(D)** Plot of projected cell spread area over time after measuring the spread area of cells shown and described in (A), with the red arrow at t = 10 minutes marking when 0.6% methylcellulose was added to the media. n = 33 for control and 32 for Arp2/3 null cells from 2 experiments for each condition.

### Viscosity-induced spreading on poly-L-Lysine-coated soft substrates generates smaller traction forces than spreading via robust Integrin-ECM engagement

In cells spreading on fibronectin, integrin-based adhesions link contractile and protrusive actin networks to the ECM(Parsons, Horwitz et al. 2010). When integrin adhesions are engaged near the protrusive leading edge of cells, they can resist the retrograde flow of treadmilling actin networks, and this generates traction forces on the substrate that drive cell spreading and shape changes. Upon para-aminoblebbistatin treatment, relieving cortical contractility that resists cell shape change might reduce the amount of traction forces required to drive cell spreading. However, the extent to which viscosity-induced spreading on poly-L-Lysine requires or influences cellular traction forces is not known. The scarce integrin ligand availability under such conditions might suggest that clutching engaged with the substrate is unlikely to produce enough of the force needed to facilitate cell flattening and spreading events. To test this, we employed time-lapse traction-force microscopy of JR20 fibroblasts plated on ∼20 kPa PDMS substrates and examined how increased viscosity-induced spreading on poly-L-Lysine compared to integrin-based spreading on fibronectin coating.

For untreated cells plated on fibronectin-coated PDMS, we measured traction forces that appeared considerably higher on traction maps than those measured for cells plated on poly-L-Lysine coated PDMS and treated with 0.6% methylcellulose (Fig.7A). Under both conditions, cells spread out significantly over the course of an hour, but while the strain energy applied to substrate does not significantly increase during viscosity-induced spreading on poly-L-Lysine, it does when cells are spreading over dense fibronectin (Fig.7B). We also found significant increases in strain energy density among cells plated on fibronectin-coated PDMS when spreading from 66% to 75% final spread area, as well as from 75% to 99%, and such significant increases were not similarly seen when cells were spreading on poly-L-Lysine following methylcellulose addition (Fig.7C). These results suggest that dense actin branching elicited by increased extracellular viscosity is sufficient to drive efficient cell spreading, even in the absence of robust adhesion and traction generation behaviors.

**Figure 7.**
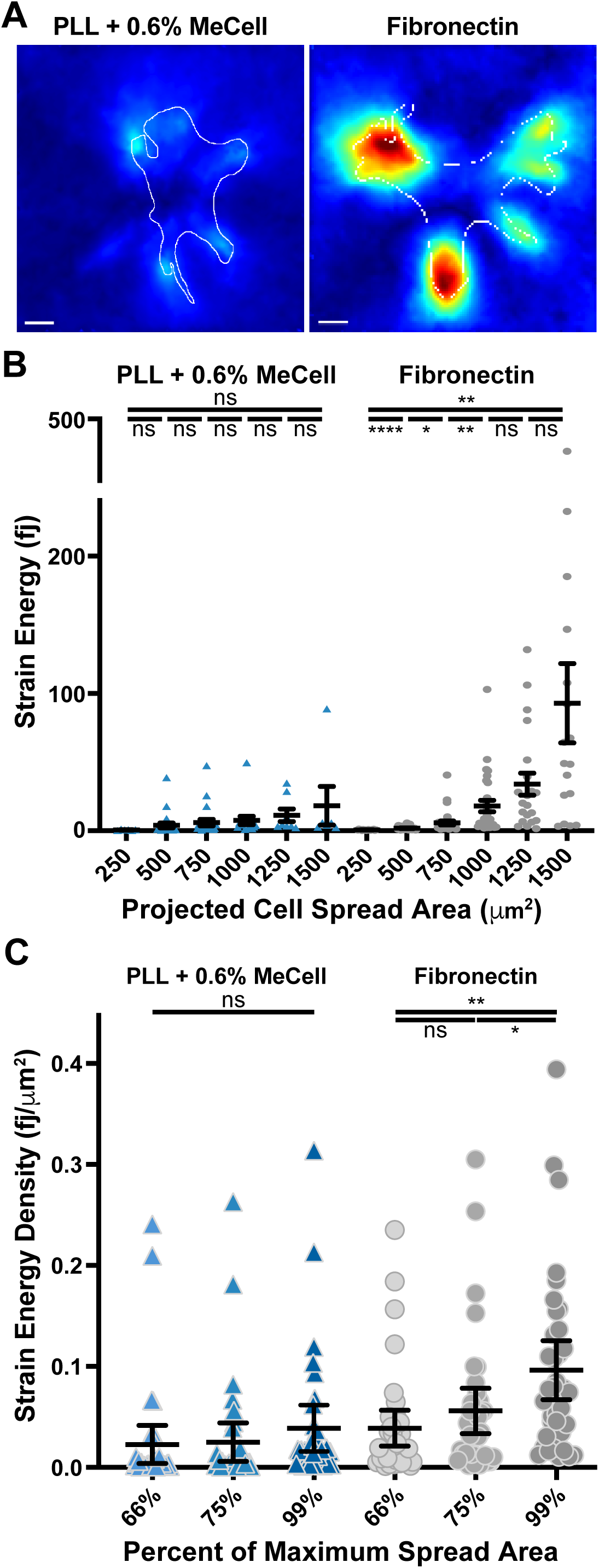
Viscosity-induced spreading on poly-L-Lysine-coated soft substrates generates smaller traction forces than spreading via robust Integrin-ECM engagement. **(A)** Traction maps generated from representative images selected from time-lapse movies of fibroblasts plated on 20 kPa PDMS substrates with red fluorescent beads attached to the surface allowing for TFM measurements and coated with either poly-L-Lysine or fibronectin. Methylcellulose was added at a final concentration of 0.6% to promote spreading on the substrates coated with poly-L-Lysine with similar dynamics to the cells plated on fibronectin. **(B)** Plot of changes in strain energy as cells spread out under the two conditions shown and detailed in (A), with error bars representing the standard error of the mean. For spread areas of 250, 500, 750, 1,000, 1,250 and 1,500 square microns, n = 6, 23, 22, 15, 8 and 6 cells for the poly-L-Lysine + 0.6%methylcellulose condition and 18, 29, 32, 29, 21 and 17 cells for the fibronectin plating condition across 3 experiments. **(C)** Plot of the strain energy density of forces applied by the same cells measured in (B) as they reached 66%, 75%, and 99% of the maximum recorded projected spread area.

### Cell protrusions generated via optogenetic Rac activation are facilitated by increased extracellular viscosity on ECM-deficient substrates

In cells spreading on fibronectin, integrin engagement and clustering provokes both protrusive signaling and mechanical resistance to retrograde actin flow. In light of our data showing that increased extracellular viscosity negates the need for integrin-based engagement for efficient cell spreading, we sought to test how signaling downstream of integrins is affected by increased extracellular viscosity conditions. Optogenetic stimulation of Rac at the leading edge of cells plated on poly-L-Lysine leads to some increased ruffling, but as we saw before (Fig.2F), little to no forward protrusion or Arp2/3-branched actin enrichment (Fig.8A-B and Movie 15). However, following treatment with methylcellulose and the subsequent spreading response, Rac activation leads to large, robust protrusions that push into regions of optogenetic stimulation (Fig.8A-B and Movie 16). Across multiple cells, Rac activation in increased extracellular viscosity conditions triggered significantly more robust protrusions when compared to when the same cell was subjected to Rac activation under pre-treatment conditions (Fig.8C). Together, these data show that cells spread in an Arp2/3-dependent manner upon an increase in extracellular viscosity, which leads to dense Arp2/3-branched actin networks in more flattened cells and protrusions, and this condition primes the leading edge to be sensitive to further protrusion induced by increased Rac activity.

**Figure 8.**
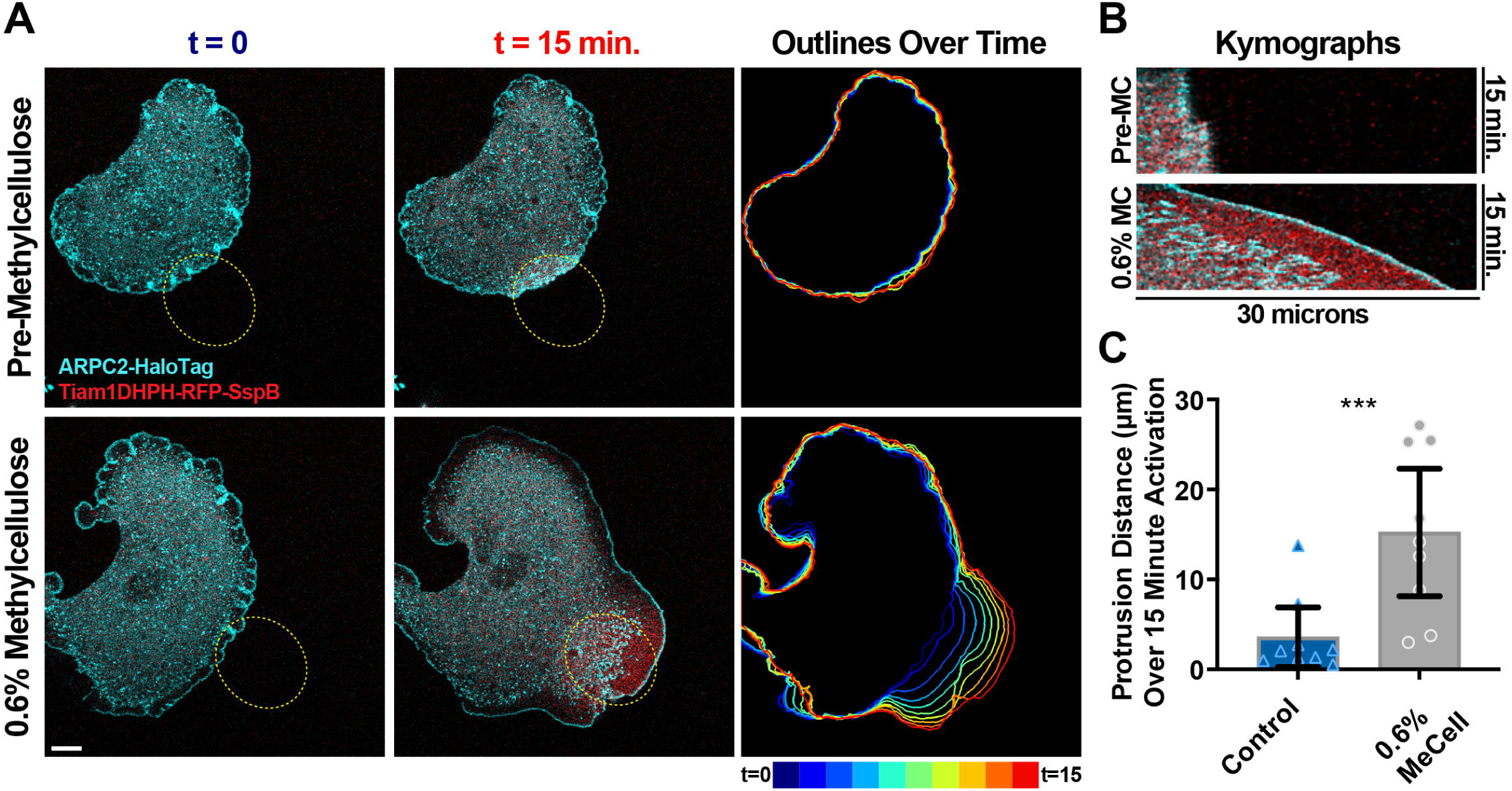
Increased extracellular viscosity facilitates cell protrusion on poly-L-Lysine during optogenetic Rac activation. **(A)** Images from representative timelapse movies of Tiam1-DH/PH-TagRFPt-SspBmicro and ARPC2-HaloTag (labeled with JF646 Halo Ligand) stably expressed by a 4HT-treated *Arpc2* knockout rescue fibroblasts that have been plated on glass coverslips coated with poly-L-Lysine and regularly stimulated with 405 nm light inside the region of interest (ROI) labeled with a yellow dashed oval starting at t = 0. These cells additionally express Venus-iLid-CAAX (not shown). The same cell is shown in all panels and was imaged during stimulation both before (top row) and after (bottom row) 0.6% methylcellulose addition to the media. Scale bar spans 10 microns. **(B)** Kymographs generated from a line (not shown) bisecting the long axis of the yellow dashed oval ROIs shown in (A). **(C)** Plot of distances the cells detailed in (A) protruded upon optogenetic activation of Rac either before or after addition of 0.6% methylcellulose to the media. n = the same 9 cells across 3 experiments for both conditions.

## DISCUSSION

In the present study, we examined the relationship between cell mechanics and signaling that control protrusive Arp2/3-branched actin organization, striving to help fill the gap in our understanding of the hierarchy or overlap between these two actin regulatory inputs. The biochemical regulation of actin dynamics through signaling from integrin adhesions has been well characterized(Romero, Le Clainche et al. 2020), with Rho-family GTPases being common downstream effectors of signaling from clustered integrin adhesion complexes. The mechanical regulation of branched actin organization has been characterized through theoretical modeling and *in vitro* experiments, and some evidence for force feedback on Arp2/3-branched actin networks in cells has been documented(Papalazarou and Machesky 2021). Most mechanical manipulations of cells have focused on altering membrane tension or examining actin network components with inferred local membrane tension fluctuations during protrusion-retraction cycles. Here, we provide a comprehensive analysis of the control of actin branch dynamics through the monitoring of endogenous Arp2/3 complexes in several different experimental contexts, and we show through both live cell imaging and genetic knockout that Arp2/3-branched actin is a key component in responding to and generating increased mechanical forces that drive cellular protrusions.

### Physical spreading alone is sufficient to enrich Arp2/3-branched actin at the periphery

Examining fibroblasts plated on surfaces coated with either fibronectin or poly-L-Lysine, which support either stronger or weaker integrin engagement respectively, revealed striking differences in branched actin organization. Integrin signaling alone is unlikely to explain these differences, as optogenetic activation of Rac does not drive protrusions in cells plated in the absence of extracellular matrix proteins(Zimmerman, Asokan et al. 2017) nor does it promote dense actin branching as we show here. One potential explanation could be that lamellipodial protrusions may require integrin signaling from nascent adhesions either beyond or in addition to activating Rac, such as PIP_3_ production downstream of FAK(Wang, An et al. 2024). However, increasing evidence of mechanical feedback on branched actin networks prompted us to investigate how aspects such as cell shape and membrane tension may play a more prominent role in dense lamellipodial actin branching.

In addition to displaying more diffuse branched actin in protrusions, fibroblasts plated for roughly one hour on poly-L-Lysine exhibit incomplete spreading, with excess membrane stored in folds, ruffles, and filopodial structures. Cells in these conditions can eventually spread out wide and flat, and in these cells, we observed more dense Arp2/3-branched actin enrichment. Global membrane tension has shown to increase through cell protrusion(Houk, Jilkine et al. 2012, De Belly, Yan et al. 2023) and cell spreading (Gauthier, Fardin et al. 2011) as well as inhibit actin polymerization and cell protrusion at elevated levels(Houk, Jilkine et al. 2012, Diz-Muñoz, Thurley et al. 2016). Thus, it is likely the shape cells adopt when efficiently spreading on dense ECM has a strong influence on branched actin density, whether through causing more confined, flattened protrusions, increasing membrane tension as cells increasingly spread out, or a likely contribution of both of these factors. Importantly, neither of these factors necessary rely upon ECM engagement specifically for dramatic changes in branched actin organization. We could not completely rule out the secretion of additional ECM molecules besides fibronectin as being important for this delayed but eventual enrichment of branched actin in cells plated on poly-L-Lysine. However, because the extent of spreading was in some cases reversible while dense Arp2/3 branching reversed along with it, this suggests that integrin engagement may not be strictly required for the dense actin branching in protrusions seen in cells plated on fibronectin but rather promotes efficient cell spreading and acquisition of a spread and flattened shape.

To further investigate this notion that cell shape following spreading on ECM, rather than the ECM specifically, has a major influence on leading edge actin organization, we promoted adhesion-independent cell spreading with manipulations that eliminated the need for traction forces to drive cell spreading. Each of the treatment conditions tested, including para-aminoblebbistatin to inhibit non-muscle myosin and cortical contractility, physical compression under agarose, sorbitol to manipulate cell volume, and methylcellulose to increase extracellular viscosity, all have their own unique caveats, but they share a similar outcome of increased branched actin enrichment in protrusions. All of these treatments ultimately result in cell spreading and/or cell flattening and suggest the resulting mechanical conditions and cell shape changes following spreading strongly promote dense actin branching in protrusions. We reason that under such conditions, increased stress between the plasma membrane and the barbed ends of actin filaments could limit accessibility of profilin-actin to sites of preferred monomer addition. This limited accessibility of actin monomers along with the functional role of capping protein(Funk, Merino et al. 2021) shift actin polymerization away from simple barbed end elongation and towards increased branching density.

### Branched actin force generation supports integrin-independent cell spreading

When plated on poly-L-Lysine, adhesions in our model fibroblast cell lines are greatly reduced or absent, and the increase in traction forces are negligible during robust spreading following increased extracellular viscosity. If not applying significantly increased traction forces to the substrate, this leads to the question of how these cells are generating the forces needed to spread and flatten upon viscosity-induced spreading on poly-L-Lysine. Branched actin is clearly required, as we showed through conditional knockout of *Arpc2*, so it is worth considering how increasingly dense branched actin might promote robust spreading in response to extracellular viscosity, if not through increased traction force generation as has been shown when cells are plated on dense fibronectin(Pittman, Iu et al. 2022).

Ruffling or buckling of the leading edge branched actin network has been linked to changes in local membrane tension at active protrusions and coincide with or control the positioning of new adhesions(Pontes, Monzo et al. 2017, Pittman, Iu et al. 2022). With increased extracellular viscosity having been shown to suppress buckling ruffles(Pittman, Iu et al. 2022) and the diminished capacity to form robust integrin adhesions in the absence of ECM, protrusions over poly-L-Lysine promoted by methylcellulose addition are likely to be generating and maintaining higher levels of local membrane tension than when produced under more standard imaging and culturing conditions. While we still do not have completely clear picture of how extracellular viscosity drives robust spreading, we propose that there may be an emphasized role for resistance to protrusive actin retrograde flow by the cytoplasmic environment, at least when cell adhesions are diminished. We would predict the physical changes that occur upon complete spreading to enhance this, including higher plasma membrane tension, an increasingly dense and more highly branched actin network that is undergoing retrograde flow, and a more axially confined lamellipodial protrusion with ruffling being suppressed. This cytoplasmic resistance to polymerizing branched actin has been previously been characterized as an important intracellular force generator in several contexts, driving processes such as endomembrane fission(Derivery, Sousa et al. 2009, Gomez and Billadeau 2009, Marchan and Bear 2025).

### Actin-membrane interface stress as a critical regulatory input to Arp2/3-branched actin

To summarize, while both cell signaling and cell mechanics are important regulatory inputs for branched actin regulation during cell spreading and protrusive migratory behaviors, our results suggest that the mechanics of actin-membrane interface stress are most influential for directly influencing branched actin architecture. We show that Rac activation is not able to increase actin branching without sufficient mechanical support, suggesting that biochemical signaling from integrin adhesions may be more important for protrusion persistence, coherence, or other regulatory behaviors that might support cell polarization, adhesion dynamics, and traction generation. In support of this model, we demonstrate that the Arp2/3-dependent response to increased extracellular viscosity was sufficient to circumvent the need for ECM to drive cell protrusion upon optogenetic Rac activation. These results are similar to those from a previous study using leukocytes that showed a necessity for integrins for 2D migration on flat surfaces but not 3D migration in more confined and restrictive environments (Lämmermann, Bader et al. 2008), and we theorize that integrin-deficient leukocytes in the restricted, 3D context would likely be more responsive upon Rac or Cdc42 activation.

It is interesting to consider other contexts besides cell spreading, migration, and protrusion where the concept of actin-membrane interface stress might apply. Branched actin has been shown to increasingly accumulate at sites of frustrated endocytosis(Wang, Galletta et al. 2016, Yang, Colosi et al. 2022), cell-cell junctions(Del Signore, Cilla et al. 2018, Efimova and Svitkina 2018, McEvoy, Sneh et al. 2022), and sites of endomembrane fission (Marchan and Bear 2025), though much less is known about the orientation of branched actin polymerization in these contexts and where, specifically, the sites of contact are at which stress is applied on the barbed ends of the branched network. In addition, it is likely that this concept has important implications throughout development and disease, though few examples have been well characterized due to the difficulty of manipulating and measuring branched actin force responsiveness *in vivo*. In one potential example, disrupting matrix metalloproteinases activity *C. elegans* embryos leads to an irregular and delayed migration of the anchor cell through the basement membrane that involves enrichment of Arp2/3-branched actin to help the cell physically force its way through (Kelley, Chi et al. 2019). Future studies will be needed to understand in more detail where and how actin-membrane interface stress and other mechanical inputs can trigger more robust Arp2/3-branch generation.

## MATERIALS AND METHODS

### Cell Culture and Cell Line Generation

Cell lines were cultured and imaged in DMEM (4.5 g/L D-Glucose, L-Glutamate, Sodium Pyruvate, Gibco, cat. no. 11995-065) supplemented with 10% FBS (MedSupply Partners) and 1x GlutaMax (Gibco, cat. no. 35050-061) at 37°C and 5% CO_2_. Cell lines used in experiments tested negative for mycoplasma using a commercial detection kit (InvivoGen). For the stable lentiviral expression of various markers (see Table S1), lentiviral production was performed using standard procedures. In brief, HEK293T cells at roughly 70% confluency in a 6 cm tissue culture dish were transfected with 500 ng each of the pLL5/7.0 expression plasmid, pCMV-VSV-G, pRSV-Rev, and pMDL-g/p plasmids using Xtremegene HP transfection reagent (Roche). Lentivirus was harvested 72 hours later by spinning down and filtering the supernatant through a 0.45 μm syringe filter before using roughly half to transduce fibroblasts freshly plated at ∼15% confluency in a 6 cm dish for 72 hours while supplemented with 4 μg/ml Polybrene. Fluorescence-positive cells with low to moderate expression levels of various markers were purified using FACS.

To induce LoxP recombination at the *Arpc2* locus, 2 μM of 4-Hydroxytamoxifen (Millipore Sigma, cat. no. H6278) was added to cultured cells at least one week before any experiments examining Arp2/3 null phenotypes. Because JR20 *Arpc2* knockouts seem to be resistant to lentiviral transduction, lentivirus generated with pLL7.0-ARPC2-HaloTag was added 24 hours after 4-Hydroxytamoxifen addition following complete media exchange in order to generate a stable ARPC2-HaloTag knockout rescue line. This timepoint allows for lentiviral transduction during a window between induced recombination at the *Arpc2* locus and complete less of *Arpc2* mRNA and protein still present for a time following recombination. Cells successfully rescued with ARPC2-HaloTag exhibit higher proliferation rates than knockouts and remained amenable to further lentiviral transduction of optogenetic components.

### Confocal Imaging and Standard Plating Conditions

For general imaging conditions, glass bottom dishes (Cellvis, cat. no. D35-20-1.5-N) were coated with 0.01% poly-L-Lysine solution (Millipore Sigma, cat. no. P4707) or Human fibronectin (Corning, cat. no. 356008) dissolved at a concentration of 10 μg/ml in PBS for 30 minutes at room temperature before washing with PBS. Fibroblasts were lifted from culture dishes using 0.25% Trypsin-EDTA (Gibco, cat. no. 25200056), and approximately 25,000 (Attofluor A-7816 imaging chambers) or 50,000-75,000 (Cellvis D35-20-1.5-N glass bottom dishes) cells were seeded for at least 30 minutes prior to imaging or fixation. Fluorescence microscopy images were captured on a Zeiss LSM800 confocal microscope fitted with a Tokai Hit stage top and Pecon large chamber incubators using a Plan-Apochromat 63X/1.4 NA Oil objective, typically at 1024x1024 pixels using 2-8x averaging and less than 0.5% of 10 mW maximum laser power for excitation. For fluorescence recovery after photobleaching (FRAP) experiments designed for visualization and quantification of actin polymerization and retrograde flow velocities, the leading-edge region was bleached (10 mW, 488 nm laser; 0.67 μs pixel dwell time; 0.76 μm pixel size; 15 iterations), and fluorescence recovery was subsequently monitored at roughly 1.5 s intervals. An example movie has been shown previously [Video S1 (Chandra, Butler et al. 2022)]. For optogenetic stimulations, 0.2% power of a 488 nm laser was used to stimulate the ROI between scans. Fixed cells for confocal imaging were prepared by treating with 4% paraformaldehyde (Electron Microscopy Services, cat. no. 15713) premixed into fresh culture media before incubation for 10 minutes at room temperature. For F-actin visualization, fixed cells were washed 3x in PBS, permeabilized with 0.2% Triton-X in PBS for 5 minutes at room temperature, and then stained with Alexa Fluor 647 Phalloidin (ThermoFisher, cat. no. A2287).

### Cell Manipulations

Agar compression was performed by casting 1% agarose dissolved in culture medium in imaging chambers (Attofluor A-7816), gently sliding the agarose puck over cells plated in glass-bottom dishes, and placing ∼8 grams of weight on top of the agarose puck during a pause in image acquisition. Additions of para-aminoblebbistatin (Cayman Chemical, cat. no. 22699), sorbitol (Sigma, cat. no. S1876), and methylcellulose (R&D systems, cat. no. HSC001) were performed by adding 1 ml total of treatment diluted in media to the 1.5 ml of media in glass-bottom imaging dishes that the cells were initially seeded into.

### Barbed End Assay

The barbed ends of free actin filaments were labeled as previously described(Bryce, Clark et al. 2005), but with slight modifications. Fibroblasts were plated into glass-bottom dishes prepared as described above and allowed to settle for 2 hours prior to labeling and fixation. Cells were washed with pre-warmed PBS and then permeabilized and labeled with 3.2 μM Alexa Fluor 488-labeled actin (Life Technologies, cat. no. A12373) in permeabilization buffer (20 mM HEPES, 138 mM KCl, 4 mM MgCl2, 3 mM EGTA, 0.2 mg/mL saponin, 1% BSA, 1 mM ATP, 3 μM phalloidin) for 30 seconds. The cells were then immediately fixed in 4% paraformaldehyde at room temperature for 10 minutes and then washed in PBS prior to imaging.

### Protein Purification and Coverslip Coating

To purify RGD peptides for coating glass coverslips, sequences coding for either Hisx6-SspB-PHSRNSGSGSGSGSGRGDNP or Hisx6-SspB-GRGDS were cloned into pQE-80L vector via Gibson Assembly, and the resulting plasmid was transformed into AVB101 competent cells (Avidity, cat. no. CVB101). Cultures were grown in autoinduction media (Novagen, cat. no. 71757) supplemented with 50 μM biotin (Sigma, cat. no. B4639) overnight at 18°C, and pelleted bacteria were frozen, thawed, resuspended and sonicated prior to protein extraction using Ni-NTA purification spin columns (ThermoFisher, cat. no. 88229) according to the manufacturer’s protocol.

For coating of glass with purified peptides, coverslips were washed with methanol and plasma cleaned for 5 minutes following drying. The coverslips were then immersed in 5% (v/v) (3-glycidyloxypropyl)trimethoxysilane (GLYMO, Sigma, cat. no. 440167) solution in methanol for 16 h at room temperature. The following day, coverslips were washed with isopropanol and heated at 105°C for one hour. Once cooled, the coverslips were coated with 1 mg/ml NeutrAvidin (ThermoFisher, cat. no. 31000) diluted in PBS overnight at room temperature in the dark. The following day, coverslips were washed with PBS and coated with 2 mg/ml of purified protein for 2 hours.

### Scanning Electron Microscopy

For scanning electron microscopy (SEM) imaging, cells where plated on round 12 mm diameter coverslips for roughly 45 minutes and fixed with 2.5% glutaraldehyde/0.15M sodium phosphate buffer (pH 7.4) for one hour at room temperature. Cells were post-fixed for 15 minutes with buffered 1% osmium tetroxide (washed 3x in water) followed by 2% tannic acid in water for 15 minutes (washed 3x in water) and then 15 minutes in aqueous 1% osmium tetroxide. The samples were then washed with three exchanges of deionized water for 10 minutes each. Samples were dehydrated in a gradient ethanol series and critical point dried using CO_2_ as the transitional solvent (Samdri-795 critical point dryer, Tousimis Research Cop., Rockville, MD). Coverslips were then mounted and sputter coated with 5 nm gold-palladium alloy (60Au:40Pd) using a Cressington 208HR Sputter Coater (Ted Pella Inc.). Images were acquired on a Zeiss Supra 25 FESEM (Carl Zeiss SMT Inc.) operating at 5 kV using a 6 mm working distance and 20 μm aperture.

### Traction Force Microscopy

Traction force microscopy was performed similarly to as previously detailed (Teo, Lim et al. 2020). In brief, a solution of compliant PDMS was prepared by mixing rigid Sylgard 184 (Dow, cat. no. 1317318) and compliant Sylgard 527 (Dow, cat. no. 1696742). Sylgard 184 was prepared at 10:1 (Part A to Part B by weight) and mixed thoroughly before degassing for 15 minutes in a tabletop vacuum chamber. While degassing, a Sylgard 527 solution was prepared at 1:1 (Part A to Part B by weight), mixed thoroughly, and degassed for 15 minutes in a tabletop vacuum chamber. Once both solutions were degassed, they were added at a 20:1 ratio (Sylgard 527 to Sylgard 184) by weight and mixed thoroughly before degassing for 15 minutes in a tabletop vacuum chamber. Once degassed, approximately 25 µL of the compliant PDMS mixture was immediately added to a plasma-cleaned #1.5 25 mm coverslip and ran for 30 seconds at 3000 RPM in a spin coater to produce an approximately 50-micron thick flat surface. The substrate was then placed in an empty pipette tip box and put into an oven set at 60°C for at least 24 hours to fully cure the PDMS. Once cured, this mixture produces substrates that are approximately 20 kPa as determined by indentation analysis(Hockenberry, Ulmer et al. 2025).

To add fluorescent beads to the traction force substrates, we first washed the cured PDMS with 100% EtOH. We then treated the surface with 2 mL of 10% APTES (Thermo, cat. no. 430941000) in 100% EtOH for 30 minutes on a shaker platform. The APTES solution was removed from the substrates, which were then washed two times with 100% EtOH before 2 mL of a 1/1000 solution of carboxylate polystyrene fluorescent beads (Thermo, cat. no. F8801) and 1 mg/mL EDC (ThermoFisher, cat. no. E7750) in diH20 was added to the substrates. Substrates were then placed on a shaker platform for one hour, the fluorescent beads and EDC solution was then removed, and the surface was washed with diH20. Substrates were either used immediately or stored for up to one week.

To add fibronectin or poly-L-Lysine and sterilize the substrates, we placed the substrates upside down in a sterile plastic 6-well dish containing 1 mL of 10 µg/mL of fibronectin (Gibco, cat. no. 33010018) or 0.1 mg/mL of Poly-L-Lysine (Millipore Sigma, cat. no. P4707) inside of a tissue culture hood. The plastic 6-well dish was then irradiated with UV light for 30 minutes to sterilize the substrates while the proteins coated the surface. Once irradiated, the substrates were washed with sterile PBS three times.

Cover glasses with traction force substrates were placed into screw-together imaging chambers (Attofluor, cat. no. A-7816), and 1 mL of culture media containing approximately 5,000 JR20 fibroblasts stably expressing GFP-Paxillin was added before a sterile plastic lid from a 35 mm plastic cell culture dish was used to cover the chamber. Cells were imaged for 1 hour, with snapshots being taken every thirty seconds. Cells plated on poly-L-Lysine were allowed to adhere for one hour before imaging for the hour following. After timelapse imaging, 100 µL of 1% SDS in DMEM was added to the substrate to remove the cells, and then a reference state image was taken of the cell-free traction bead field.

### Traction force analysis

Traction force analysis was performed using μ-inferforce, a MATLAB TFM package (Han, Oak et al. 2015), as previous described (King, Butler et al. 2022). Briefly, images were drift corrected through Efficient Subpixel Registration before displacement fields were calculated grid points and PIV suite. The template size was set to 11 pixels, and maximum displacement was set to 20 pixels. Displacement fields were corrected through filtering vector field outliers with a normalized displacement residual of 2. Force fields were constructed using FTTC, a young’s modulus value of 20, and a gel thickness of 50. A constant regularization parameter of 1e^-5^ was used for all images.

Traction maps were analyzed by masking the image with a 5-micron dilated cell boundary determined from the GFP-Paxillin channel. Masks of the cell were obtained by manually segmenting the cell boundary from the GFP-Paxillin channel using the Fiji distribution of ImageJ(Schindelin, Arganda-Carreras et al. 2012). The values for the traction magnitude and strain energy were computed as previously described(Butler, Tolić-Nørrelykke et al. 2002).

### Image quantification and presentation

The Fiji distribution of ImageJ(Schindelin, Arganda-Carreras et al. 2012) was used to measure areas, rates, and intensities from imaging datasets. Mean intensities along cell edges were measured with a freehand line roughly 1 micron wide at and within the cell edge of the labeled regions of the largest cell protrusion. Line scans were drawn perpendicular from the edge near the middle of the largest protrusion inward, avoiding regions with obvious ruffles and folds to try to capture regions of flat, even lamellipodia. Retrograde flow rates were measured from slopes of recovering signal in kymographs generated following bleaching of GFP-β-actin at the edge of lamellipodial protrusions. These kymographs were generated from a rectangular region approximately 5 μm long and 1 μm wide, centered near the edge of the protrusion, and oriented longwise along the direction of retrograde flow. Velocities were calculated as distance/time, with the distance being the width of the base of the triangle in the resulting kymograph generated this way, and the time being the number of frames (lines) from the base to the top point at t =0 post-bleach in the kymograph. An annotated example has been shown previously [Figure 5 (Chandra, Butler et al. 2022)], with the retrograde flow rate reported here being analogous to the “polymerization rate” detailed previously, both referring to the total recovered actin network distance. The density of GFP-Paxillin foci was determined by manually counting foci that appeared roughly 100-300 nm in diameter within ∼2 microns along the largest protrusive edge. The “detect particles” tool was used following manual thresholding for cell segmentation, and mean cell intensities of segmented cells and subsequent measures were collected using macros to reduce the ROI size in one pixel steps before being used to calculate whole-cell Full Width Half Max and Max/Mean values. Bleach correction for ARPC2-mScarlet line scans at the leading edge was performed by normalizing to the background intensity regions around the nucleus prior to quantification of ARPC2-mScarlet distribution before and after cell compression under agarose pucks.

Raw images were copied from Fiji, and the Despeckle, Gaussian Blur (Radius: 0.5 pixels), and Unsharp Mask filters were applied in Photoshop (Adobe) before assembling figures using Illustrator (Adobe). Brightness and contrast were also adjusted in Photoshop and similarly applied to both control and experiment images across datasets. Prism (GraphPad) was used to plot data with 95% Confidence Interval (unless otherwise noted in the figure legend) and to perform statistical analyses (unpaired t tests with Welch’s correction). Significance notations in figures represent p ≤ 0.05 as *, p ≤ 0.01 as **, p ≤ 0.001 as ***, and p ≤ 0.0001 as ****.

## MATERIALS AVAILABILITY

All raw data, spreadsheets containing measurements, scripts, novel cell lines, and plasmids are available upon reasonable request.

## ACKNOWLEDGMENTS

We would like to thank Jason Haugh, Richard Cheney, and Dyche Mullins for valuable feedback and discussions. We would also like to thank Mark Hazelbaker for assistance with image quantification macros. We would like to give a special thanks to William Polacheck and McKenzie Garcia for insightful conversations as well their efforts towards a better understanding and interpretation of the effects of extracellular viscosity manipulations. We would like to thank Kristen White and Jillann Madren in the UNC Microscopy Services Laboratory for their help collecting SEM data. The Microscopy Services Laboratory, Department of Pathology and Laboratory Medicine, is supported in part by P30 CA016086 Cancer Center Core Support Grant to the UNC Lineberger Comprehensive Cancer Center. Research reported here was supported by funding awards F32GM131578 (M.T.B.), R35GM158040 (W.R.L.), and R35GM130312 (J.E.B.) from the National Institute of General Medical Sciences, and W.R.L. acknowledges additional support from the Packard Fellowship in Science and Engineering. The content is solely the responsibility of the authors and does not necessarily represent the official views of the National Institutes of Health.

## AUTHOR CONTRIBUTIONS

M.T.B, M.A.H., J.E.B., and W.R.L designed research. M.T.B. and M.A.H. performed research and analyzed data. M.T.B., M.A.H., and H.H.T. contributed to data visualization and presentation. M.T.B. and J.E.B. assembled and wrote the manuscript.

---

**Figure 1 Supplement 1. Characterization of endogenous *Arpc2-mScarlet* knock-in lines. (A)** Image showing the resulting PCR products following amplification of genomic DNA isolated from the indicated JR20 Fibroblast CRISPR-modified clones and separation via gel electrophoresis. The single band product observed for Clone 48 was confirmed to be a clean *Arpc2-mScarlet* knock-in through subsequent Sanger sequencing. **(B)** Representative confocal images of groups of cells predicted to be either monoallelic (Clone 18 and Clone 20) or biallelic (Clone 46 and Clone 48) *Arpc2-mScarlet* knock-ins plated in 10 μg/ml fibronectin-coated glass-bottom imaging dishes. Images are of the same scale. **(C)** Graphs of values from traced whole cell or leading edge measurements of cells plated as shown in (B). Error bars represent the 95% confidence interval.

**Figure 1 Supplement 2. Barbed end density increases with ARPC2-mScarlet enrichment in fibroblasts plated on surfaces coated with dense fibronectin ECM. (A)** Images of fibroblasts endogenously expressing ARPC2-mScarlet after simultaneously pulsing labeled Alexa Fluor 488 Actin and unlabeled phalloidin under conditions that make the cells slightly permeable to label the barbed ends of stabilized actin filaments. Scale bar spans 10 microns. **(B)** Plot of mean intensity of Alexa Fluor 488 Actin in cells shown and described in (A). n = 9 cells for each condition. **(C)** Plot of values for mean edge intensity of pulsed labeled Alexa Fluor 488 Actin shown in (B) against the similarly measured ARPC2-mScarlet in the same cell, with linear regression lines of similar slopes demonstrating a comparable relationship between barbed end and ARPC2-mScarlet densities in cells plated on poly-L-Lysine (blue dots) when compared to fibronectin (grey dots).

**Figure 1 Supplement 3. Branched actin enrichment in fibroblasts plated on fibronectin depends on Integrin engagement. (A)** Images of endogenous ARPC2-mScarlet and exogenous LifeAct-miRFP670 and GFP-Paxillin stably expressed by fibroblasts plated on surfaces that have been coated with either poly-L-Lysine or fibronectin. Red arrows mark the edge of the largest cell protrusion where the presence or absence of small paxillin clusters can be seen. Scale bar spans 5 microns. **(B)** Plot of density of GFP-Paxillin clusters around the edge of the largest cell protrusion under the conditions shown in (A). n = 16 cells from 3 experiments. Note that the data from cells on poly-L-Lysine coated glass in panels (A & B) were collected at the same time as other experimental conditions for a separate study, and the images and values for the fibronectin condition here have been published as the 10 μg/ml control condition previously(Chandra, Butler et al. 2022). **(C)** Images of endogenous ARPC2-mScarlet and exogenous LifeAct-miRFP670 and GFP-Paxillin stably expressed by fibroblasts plated on surfaces that have been coated with covalently-linked poly-L-Lysine, fibronectin, purified GRGDS peptides, or purified PHSRNGRGDNP peptides. Scale bar spans 5 microns. **(D)** Plot of mean ARPC2-mScarlet intensities at and within roughly 1 micron of the cell edge along the largest protrusion in fibroblasts plated as described and shown in (C). n = 29, 27, 14 and 38 for poly-L-Lysine, fibronectin, GRGDS, and GRGDNP coating conditions, respectively.

**Figure 3 Supplement 1. Fibronectin promotes efficient spreading and flattening of fibroblasts. (A-B)** Scanning electron microscopy (SEM) images of fibroblasts plated on glass coverslips coated with poly-L-Lysine or fibronectin. The panel on the right is an enlarged view of the boxed region in the panel to the left. **(C)** Plot of projected cell spread area in fibroblasts plated on either fibronectin or poly-L-Lysine over time. **(D)** Sample images and accompanying line scans measured from regions marked with yellow-shaded regions and arrows for cells plated on poly-L-Lysine. Note the maintained increase in both spread area and ARPC2-mScarlet density shown in the panels for the top section (i) and the increase and decrease of ARPC2-mScarlet density that is seen upon the increase and decrease of cell spread area shown in the panels for the bottom section (**ii)**. Scale bars span 10 microns.

**Figure 3 Supplement 2. Segmentation and quantification of Arp2/3 distribution near the cell periphery during cell spreading.** Representative images of ARPC2-mScarlet endogenously expressed by fibroblasts plated on glass coverslips coated with poly-L-Lysine as they a projected cell spread area that is either 66% (top) or 99% (bottom) of the maximum spread area recorded. Yellow lines show boundaries of segmentation of the example cell around the outer edge and 5 microns into the cell from the edge in 1-pixel steps, which was used to calculate the F.W.H.M. and maximum/mean intensities as measures of branched actin density and distribution around the cell periphery during spreading shown in Figure 3.

**Figure 4 Supplement 1. Physically flattening cells or manipulating cell volume with osmotic pressure leads to enriched protrusive branched actin. (A)** Frames from a representative time lapse movie of endogenous ARPC2-mScarlet expressed by fibroblasts plated on glass coverslips coated with poly-L-Lysine and compressed under weighted agarose pucks at t = 5 minutes to physically force cell flattening and spreading. Scale bar spans 10 microns. **(B)** Plot of the fold change in projected cell spread area after physically compressing cells at t = 10 minutes (red arrow) as shown in (A). n = 25 cells from 2 experiments. **(C)** Plot of ARPC2-mScarlet intensities along line scans starting near the middle of the largest cellular protrusion, aligned perpendicular to the cell edge, and directed inwards while avoiding regions of obvious ruffles and folds in cells plated as detailed in (A-B) following bleach correction pre-(t = 0) and post-compression (t = 60 minutes). **(D)** Frames from a time lapse movie of endogenous ARPC2-mScarlet expressed by fibroblasts plated on glass coverslips coated with poly-L-Lysine and treated with sorbitol at a final concentration of 0.25M at t = 10 minutes. Scale bar spans 10 microns. **(E)** Plot of the fold change in projected cell spread area after treating with 0.25M sorbitol at t = 10 minutes (red arrow) as shown in (D). n = 32 cells from 2 experiments. **(F)** Plot of ARPC2-mScarlet intensities along line scans starting near the middle of the largest cellular protrusion, aligned perpendicular to the cell edge, and directed inwards while avoiding regions of obvious ruffles and folds in cells plated as detailed in (D-E) before (t = 0) and after (t = 90 minutes) sorbitol addition. **(G)** Plot of mean ARPC2-mScarlet intensity around and within ∼1 micron of the edge of the largest cell protrusion in cells shown and detailed in (D-F) before (t = 0) and after (t = 90 minutes) sorbitol addition. **(H)** Plot of Full Width Half Max values calculated from the line scans shown in (F). **(I)** Representative scanning electron microscopy (SEM) images of fibroblasts plated on a glass coverslip coated with poly-L-Lysine and treated with 50 μM para-aminoblebbistatin. Right panel is an enlarged view of the boxed region in the panel to the left. **(J)** Representative scanning electron microscopy (SEM) images of fibroblasts plated on a glass coverslip coated with poly-L-Lysine and treated with 0.25M sorbitol. Right panel is an enlarged view of the boxed region in the panel to the left.

**Figure 6 Supplement 1. Neither wild type nor Arp2/3 null fibroblasts plated on dense fibronectin ECM exhibit robust cell spreading in response to increased extracellular viscosity. (A)** Frames from representative time lapse movies of endogenous ARPC2-mScarlet expressed by control cells and exogenous GFP-Paxillin and LifeAct-miRFP670 stably expressed by both control and 4-HT-treated *Arpc2* knockout fibroblasts plated on glass coverslips coated with fibronectin and treated with 0.6% methylcellulose at t = 10 minutes. Scale bars span 10 microns. **(B)** Plot of projected cell spread area over time measured among cells shown and described in (A), with the red arrow at t = 10 minutes marking when 0.6% methylcellulose was added to the media. n = 27 for control and 15 for Arp2/3 null cells from 2 experiments.

**Table S1.** Cell lines used in this study.

| Cell Line / Background | Notes / Labels (promoters in parenthesis) | Figures Used |
| --- | --- | --- |
| JR20 Fibroblasts<br>(Parental Line) | Parental JR20 fibroblasts immortalized via *Ink4a/Arf*<sup>-/-</sup> derived from mouse detailed in Rotty *et al.* 2015 *Dev. Cell*.<br>Stable/Lentiviral pLL5.0-GFP-Paxillin (MSCV 5' LTR) | 7 |
| JR20 Fibroblasts<br>Endogenous *Arpc2-mScarlet*<br>(various clones) | Methods for *Arpc2* C-terminus-mScarlet CRISPR knockin detailed in Chandra *et al.* 2022 *Biophysical Journal* | 1S1 |
| JR20 Fibroblasts<br>Endogenous *Arpc2-mScarlet*<br>(Clone 48)<br>*Fn1* (Fibronectin) CRISPR-mediated knockout (Clone 14) | *Fn1*<sup>-/-</sup> Clone 14 alleles: 1 bp & 455 bp insertions (both causing frame shifts) near N-terminus (following V22)<br>Stable expression of the following when indicated:<br>Lentiviral pLL5.0-GFP-β-Actin (MSCV 5' LTR)<br>Lentiviral pLL5.0-GFP-Paxillin (MSCV 5' LTR)<br>Lentiviral pLL5.0-LifeAct-miRFP670 (MSCV 5' LTR)<br>Lentiviral pLL5.0-mEmerald-hWAVE1 (MSCV 5' LTR)<br>Lentiviral pLL5.0-Cortactin-mEmerald (MSCV 5' LTR) | 1, 1S2, 1S3, 2, 3, 3S1, 3S2, 4, 4S1, 5, 6, 6S1 |
| JR20 Fibroblasts<br>Endogenous *Arpc2-mScarlet*<br>(Clone 48)<br>*Fn1* (Fibronectin) CRISPR-mediated knockout (Clone 14)<br>4-Hydroxytamoxifen (4HT) added to induce conditional loss of *Arpc2-mScarlet* expression | Stable expression, transduced prior to 4-HT treatment:<br>Lentiviral pLL5.0-GFP-Paxillin (MSCV 5' LTR)<br>Lentiviral pLL5.0-LifeAct-miRFP670 (MSCV 5' LTR) | 6 & 6S1 |
| JR20 Fibroblasts<br>4-Hydroxytamoxifen (4HT) added to induce conditional loss of *Arpc2* expression<br>Stable/Lentiviral Rescue with pLL7.0-ARPC2-HaloTag (CMV) | Stable expression of iLid Optogenetics tools detailed in Guntas *et al.* 2014 *PNAS*:<br>Lentiviral pLL7.0-Venus-iLid-CAAX (CMV)<br>Lentiviral pLL7.0-Tiam1(64-437)-TagRFPt-SspBμ (CMV) | 2 & 8 |

## SUPPLEMENTAL MOVIE LEGENDS

**Supplemental Movie 1 for Figure 1A**.

Representative timelapse movie of endogenous ARPC2-mScarlet expressed by fibroblasts plated on glass coated with poly-L-Lysine. Scale bar spans 10 microns, and the time stamp shows MM:SS elapsed.

**Supplemental Movie 2 for Figure 1A**.

Representative timelapse movie of endogenous ARPC2-mScarlet expressed by fibroblasts plated on glass coated with fibronectin. Scale bar spans 10 microns, and the time stamp shows MM:SS elapsed.

**Supplemental Movie 3 for Figure 2A**.

Representative timelapse movie of fibroblasts plated on glass coated with poly-L-Lysine while stably expressing mEmerald-WAVE1 shown both alone (left) and merged (green) with endogenous ARPC2-mScarlet (magenta). Scale bar spans 10 microns, and the time stamp shows MM:SS elapsed.

**Supplemental Movie 4 for Figure 2A**.

Representative timelapse movie of fibroblasts plated on glass coated with fibronectin while stably expressing mEmerald-WAVE1 shown both alone (left) and merged (green) with endogenous ARPC2-mScarlet (magenta). Scale bar spans 10 microns, and the time stamp shows MM:SS elapsed.

**Supplemental Movie 5 for Figure 2F**.

Representative timelapse movie of Tiam1-DH/PH-TagRFPt-SspBmicro (left) and ARPC2-HaloTag visualized with JF646 Halo Ligand (right) stably expressed by 4HT-treated *Arpc2* knockout fibroblasts that have been plated on glass coverslips coated with poly-L-Lysine. The yellow ROI shown on the first frame marks where stimulation with 405 nm light occurred between each frame, beginning between frames taken at t = 00:13 and 00:17 (time stamp shows MM:SS elapsed). See Figure 2E for scale.

**Supplemental Movie 6 for Figure 2F**.

Representative timelapse movie of Tiam1-DH/PH-TagRFPt-SspBmicro (left) and ARPC2-HaloTag visualized with JF646 Halo Ligand (right) stably expressed by 4HT-treated *Arpc2* knockout fibroblasts that have been plated on glass coverslips coated with fibronectin. The yellow ROI shown on the first frame marks where stimulation with 405 nm light occurred between each frame, beginning between frames taken at t = 00:13 and 00:17 (time stamp shows MM:SS elapsed). See Figure 2E for scale.

**Supplemental Movie 7 for Figure 3A and Figure 3 Supplement 1Dii.**

Representative timelapse movie of endogenous ARPC2-mScarlet expressed by fibroblasts plated on glass coverslips coated with poly-L-Lysine. Scale bar spans 10 microns, and the time stamp shows MM:SS elapsed (starts an additional 30 minutes later relative to Supplemental Movie 8).

**Supplemental Movie 8 for Figure 3A**.

Representative timelapse movie of endogenous ARPC2-mScarlet expressed by fibroblasts plated on glass coverslips coated with fibronectin. Scale bar spans 10 microns, and the time stamp shows MM:SS elapsed (starts 30 minutes sooner after plating relative to Supplemental Movie 7).

**Supplemental Movie 9 for Figure 4A**.

Representative timelapse movie of fibroblasts plated on glass coverslips coated with poly-L-Lysine while expressing endogenous ARPC2-mScarlet (magenta) and exogenous GFP-Paxillin (green) that were treated with an equal volume of DMSO used for para-aminoblebbistatin treatments, used as a control for such treatments, between frames 1 and 2. Scale bar spans 10 microns, and the time stamp shows MM:SS elapsed.

**Supplemental Movie 10 for Figure 4A**.

Representative timelapse movie of fibroblasts plated on glass coverslips coated with poly-L-Lysine while expressing endogenous ARPC2-mScarlet (magenta) and exogenous GFP-Paxillin (green) that were treated with 50 μM para-aminoblebbistatin at t = 10:00. Scale bar spans 10 microns, and the time stamp shows MM:SS elapsed.

**Supplemental Movie 11 for Figure 4 Supplement 1A.**

Representative timelapse movie of fibroblasts plated on glass coverslips coated with poly-L-Lysine while expressing endogenous ARPC2-mScarlet that were compressed under weighted agarose pucks applied at t = 5:00. Scale bar spans 10 microns, and the time stamp shows MM:SS elapsed.

**Supplemental Movie 12 for Figure 4 Supplement 1D.**

Representative timelapse movie of fibroblasts plated on glass coverslips coated with poly-L-Lysine while expressing endogenous ARPC2-mScarlet (magenta) and exogenous GFP-Paxillin (green) that were treated with 0.25M Sorbitol at t = 10:00. Scale bar spans 10 microns, and the time stamp shows MM:SS elapsed.

**Supplemental Movie 13 for Figure 5A**.

Representative timelapse movie of fibroblasts plated on glass coverslips coated with poly-L-Lysine whiole expressing endogenous ARPC2-mScarlet (magenta) and exogenous GFP-Paxillin (green) that were left unperturbed as controls for methylcellulose treatments. Scale bar spans 10 microns, and the time stamp shows MM:SS elapsed.

**Supplemental Movie 14 for Figure 5A**.

Representative timelapse movie of fibroblasts plated on glass coverslips coated with poly-L-Lysine while expressing endogenous ARPC2-mScarlet (magenta) and exogenous GFP-Paxillin (green) that were treated with a final concentration of 0.6% methylcellulose to increase extracellular viscosity at t = 10:00. Scale bar spans 10 microns, and the time stamp shows MM:SS elapsed.

**Supplemental Movie 15 for Figure 8A**.

Representative timelapse movie of Tiam1-DH/PH-TagRFPt-SspBmicro (red) merged with ARPC2-HaloTag visualized with JF646 Halo Ligand (cyan) stably expressed by 4HT-treated *Arpc2* knockout fibroblasts that have been plated on glass coverslips coated with poly-L-Lysine and stimulated in standard culture media. The yellow ROI shown on the first frame marks where stimulation with 405 nm light occurred between each frame, beginning between frames taken at t = 00:24 and 00:30. Scale bar spans 10 microns, and the time stamp shows MM:SS elapsed.

**Supplemental Movie 16 for Figure 8A**.

Representative timelapse movie of Tiam1-DH/PH-TagRFPt-SspBmicro (red) merged with ARPC2-HaloTag visualized with JF646 Halo Ligand (cyan) stably expressed by 4HT-treated *Arpc2* knockout fibroblasts that have been plated on glass coverslips coated with poly-L-Lysine and stimulated in media containing 0.6% methylcellulose (Note that this is the same cell shown in Movie 15). The yellow ROI shown on the first frame marks where stimulation with 405 nm light occurred between each frame, beginning between frames taken at t = 00:24 and 00:30. Scale bar spans 10 microns, and the time stamp shows MM:SS elapsed.

