## Supplementary figures and images for "Actin-membrane interface stress regulates Arp2/3-branched actin density during lamellipodial protrusion"

### Sup. Fig. 1-1

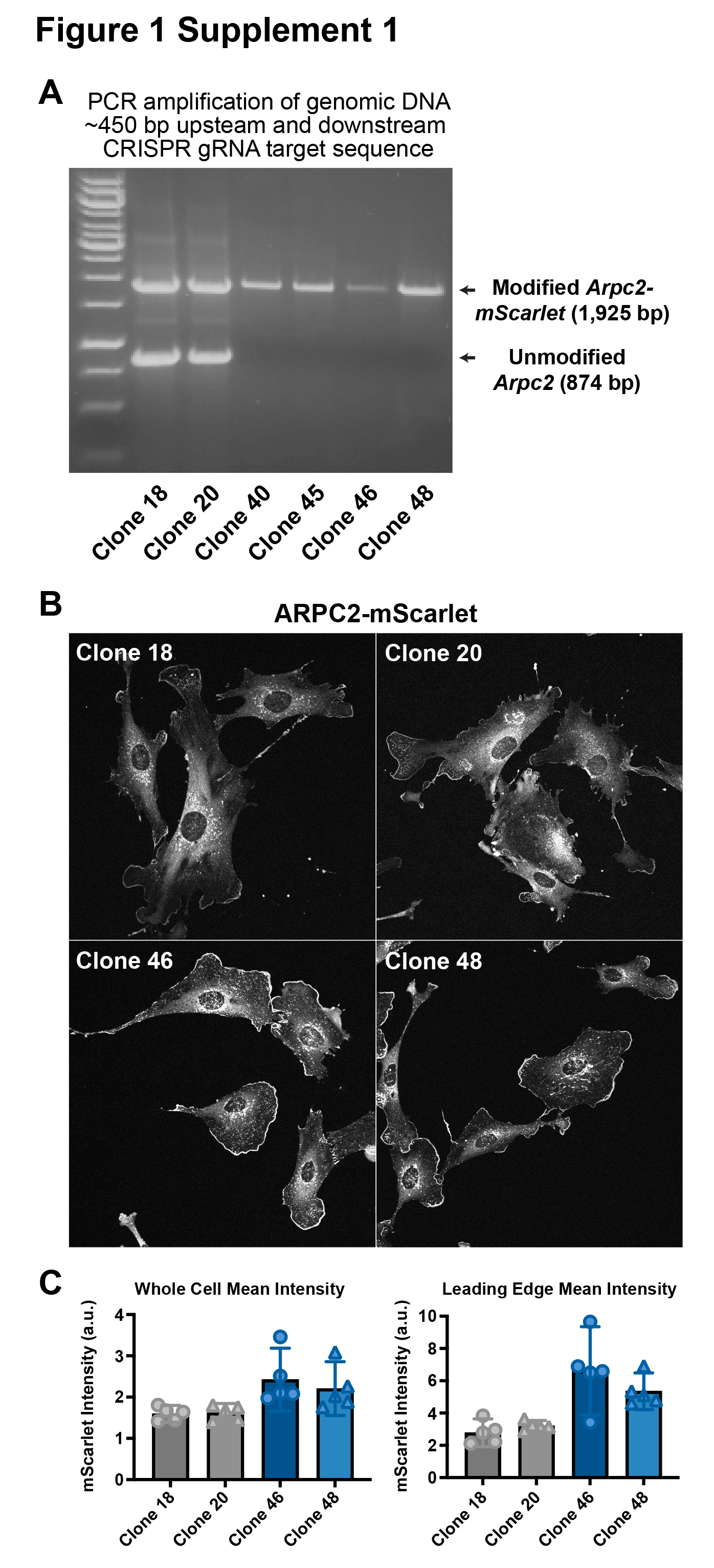

### Sup. Fig. 1-2

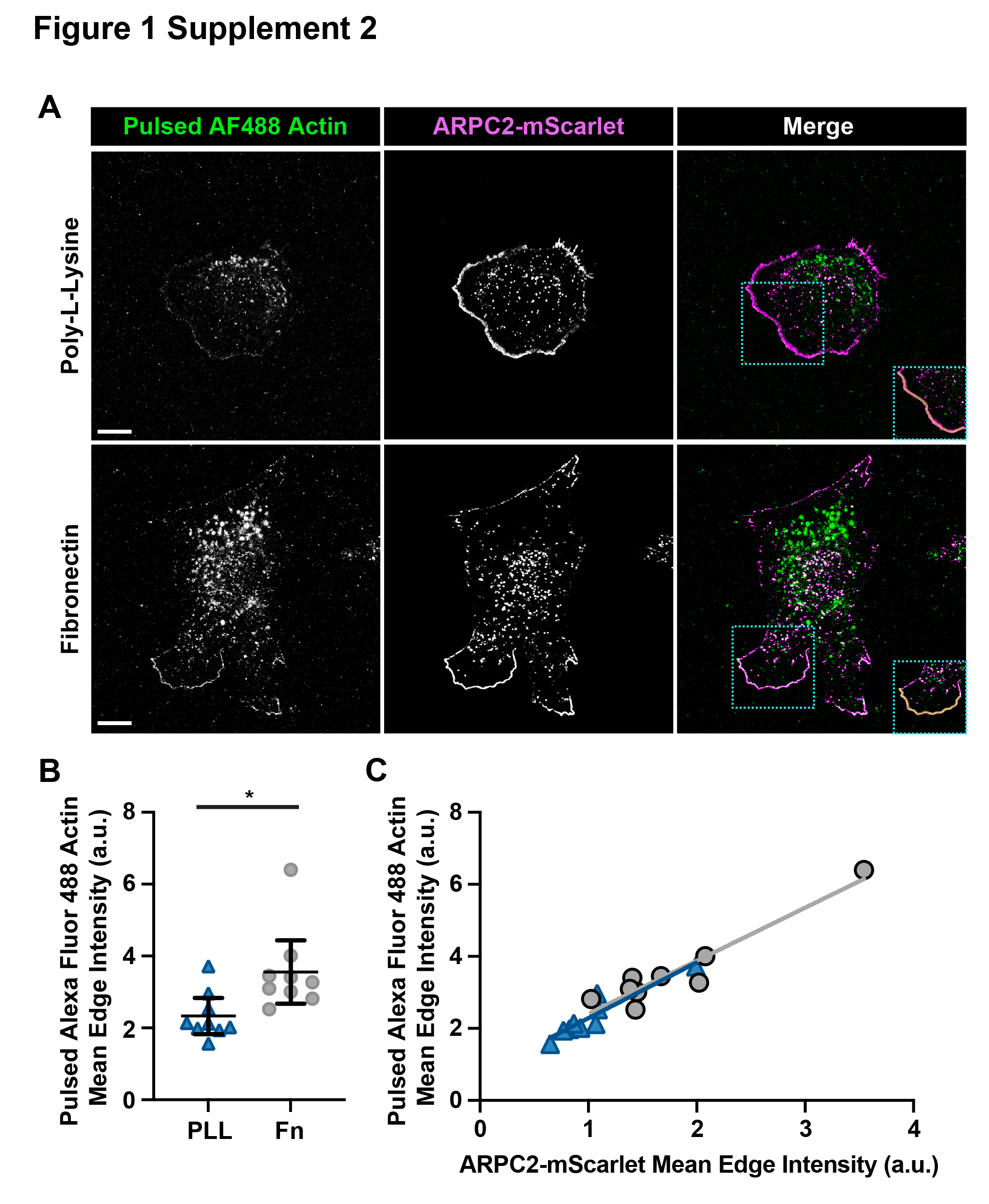

### Sup. Fig. 1-3

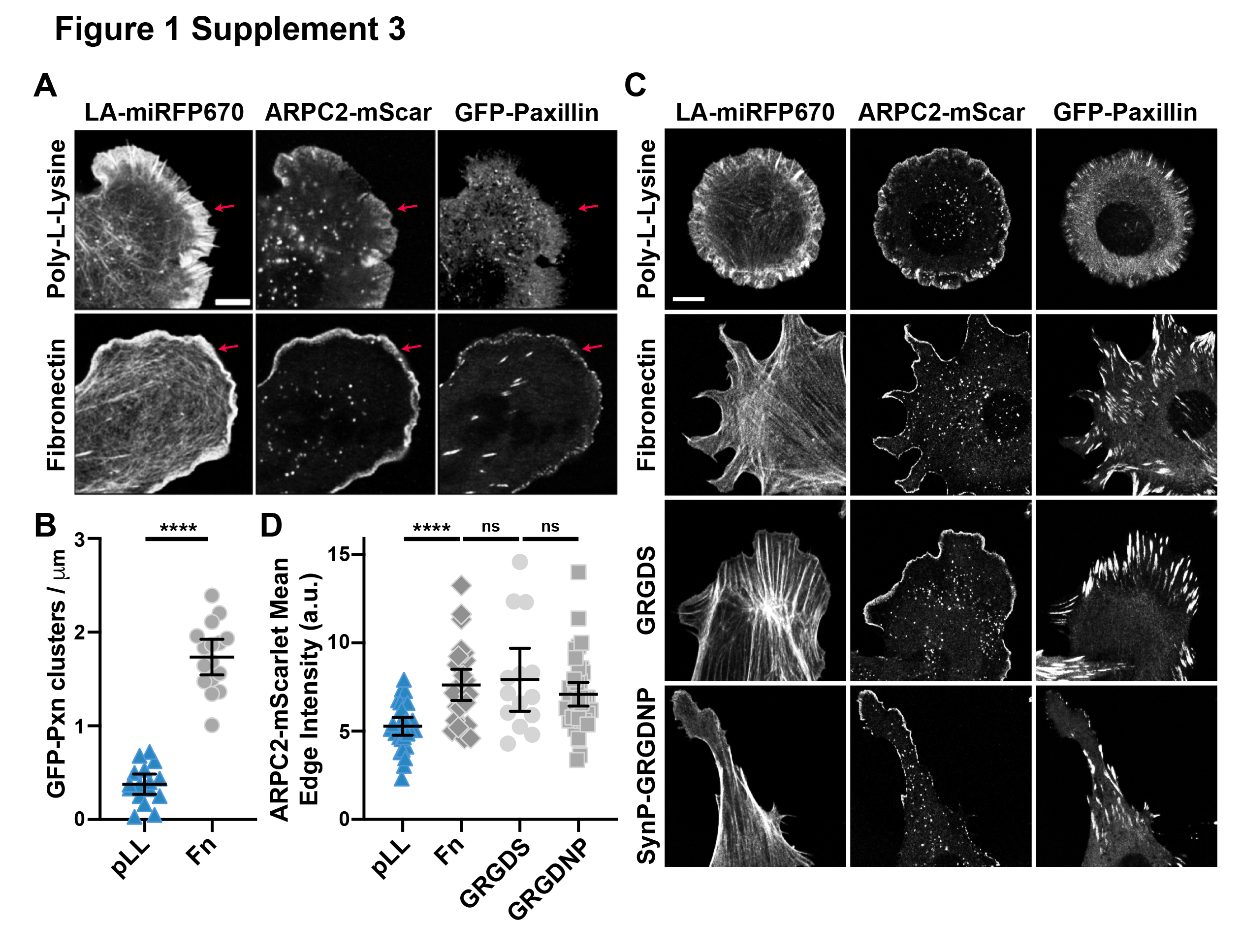

### Sup. Fig. 3-1

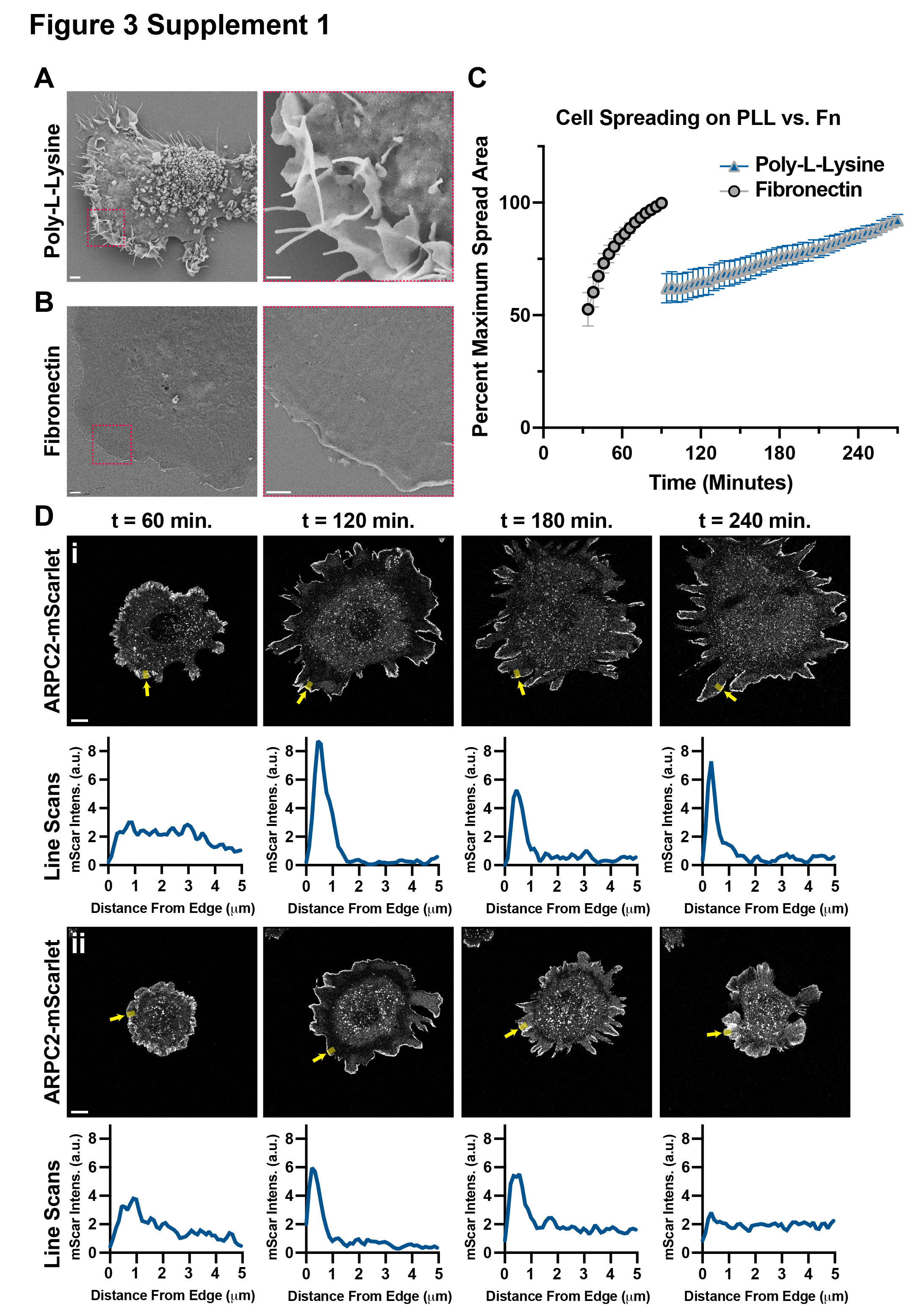

### Sup. Fig. 3-2

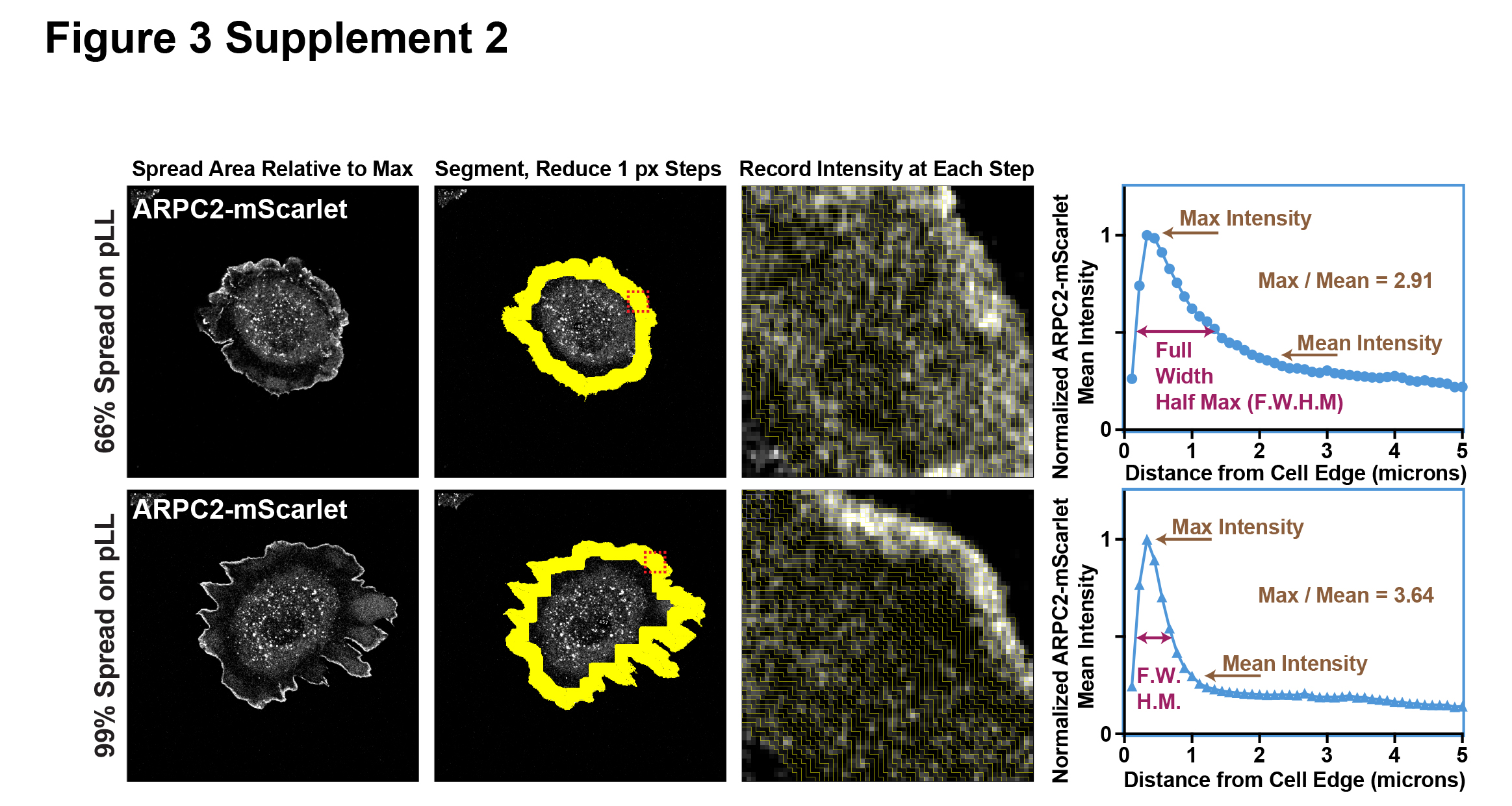

### Sup. Fig. 4

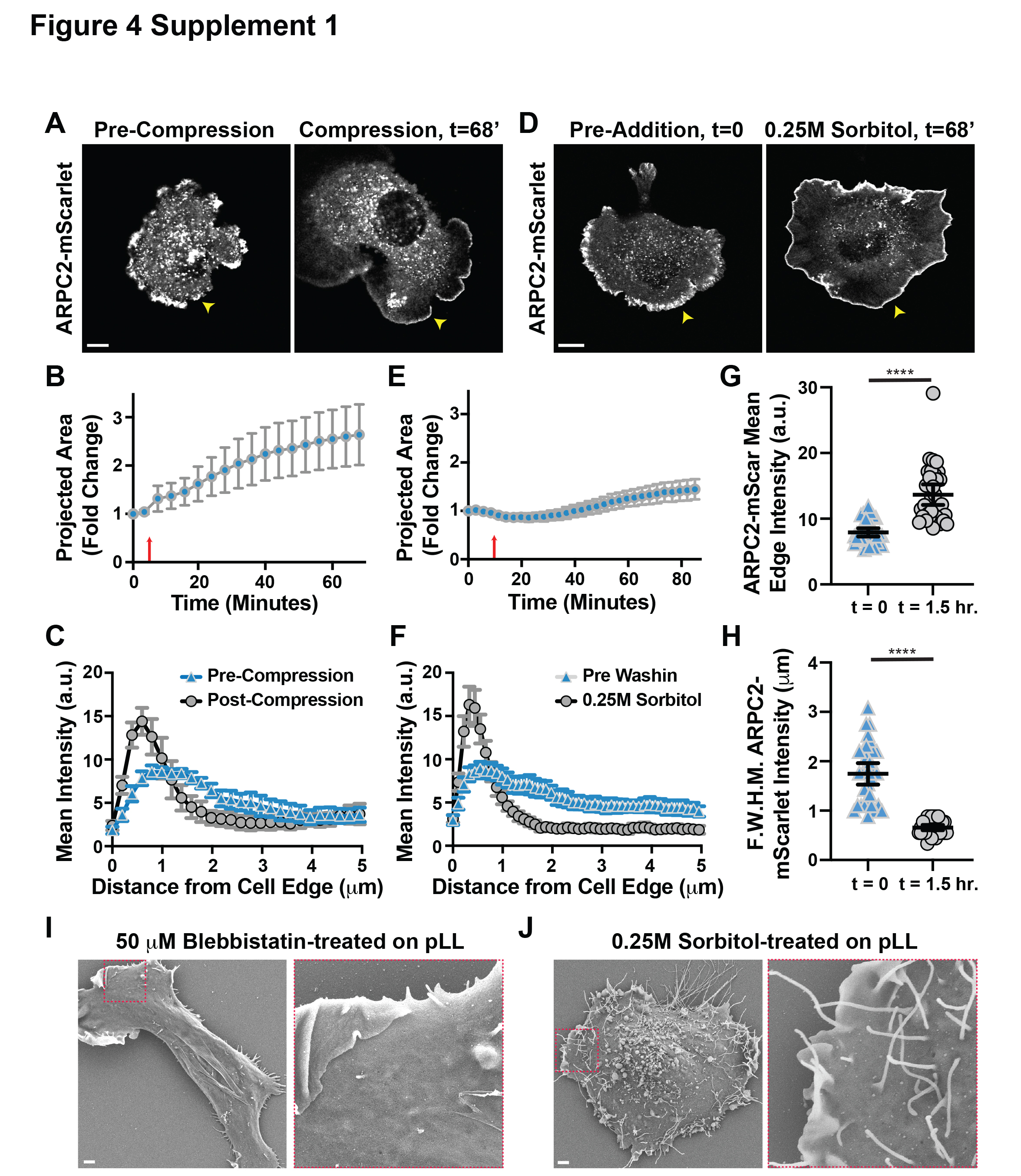

### Sup. Fig. 6

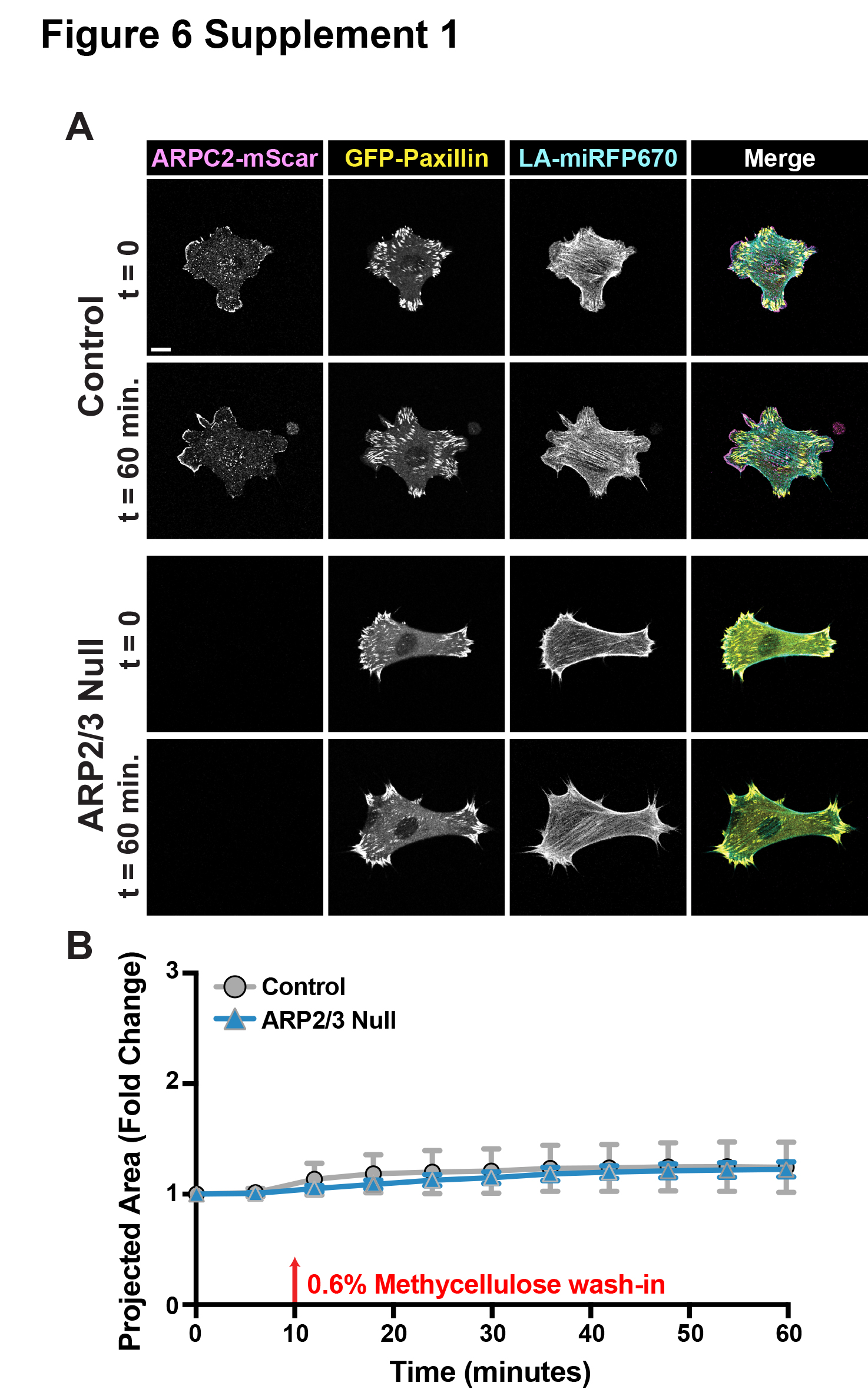
